# Role of RIGI, MDA5 and interferon alpha of duck in Duck Plague infection – a novel report

**DOI:** 10.1101/2022.01.26.477779

**Authors:** Subhomoy Pal, Samiddha Banerjee, Abantika Pal, Dhruba Jyoti Kalita, Subhasis Batabyal, Manti Debnath, Argha Chakraborty, Gourhari Mondal, Barun Roy, Paresh Nath Chatterjee, Jayanta Kumar Chatterjee, Aruna Pal

## Abstract

Duck Plague (DP) or Duck viral enteritis is an acute contagious and highly fatal disease in water fowl commonly caused by Anatidalphavirus-1 belonging from *Herpesviridae* family and contains double stranded DNA as genetic material. Pathogen associated molecular pattern (PAMP)s when identified by Pathogen Recognition Receptor (PRR)s acts as effective immunity system action against the pathogen. Melanoma Differentiation-Associated protein 5 (MDA5) and Retionic Acid Inducible Gene I (RIG1) are protein sensor commonly sense for viral double stranded RNA and helps for pro-inflammatory cytokine expression. Gut Associated Lymphoid Tissue (GALT)s have important role in immune response. The current study depicts the important role of three important immune response genes as RIGI, MDA5 and INFalpha in duck plague infestation for the first time. *In silico* studies followed by differential mRNA expression of RIG1, MDA5 and INFalpha was employed to detect effectiveness of gut associated immune responsiveness in liver, where kupfer cells are the major immune response cells. This was further confirmed through histological section of liver, kupfer cell and immunohistochemistry. This will be helpful to identify molecular mechanism of host innate immunity through duck plague virus infection in indigenous duck. This information may be helful for production of duck with the inherent resistance against duck plague virus infection through suitable biotechnological approaches as gene editing.Due to this inherent nature of better immunity in terms of resistance to other common avian diseases, duck will evolve as one of the major sustainable poultry species.The current study explores the scope to study host immunity against herpes virus in animal model.

## Introduction

Duck virus enteritis (DVE) also known as Duck Plague (DP) is one of the most common diseases in duck caused by Anatid alphaherpesvirus-1 or DVE virus of the family *Herpesviridae.* The disease causing agent is characterized as enveloped, double stranded DNA (dsDNA) virus (**Li et al., 2016**). It is an acute contagious highly fatal disease causing a high mortality percentage among duck population along with decrease egg production resulting in huge economic loss. (**You et al., 2018**)

The innate immune system is known as evolutionary defense strategy basically the front line defense system against the pathogenic microbes and boost up the adaptive immunity. It is activated when pathogen associated molecular patterns (PAMPs) is identified by pathogen recognition receptors (PRRs) which then leads to the production of interferon and cytokines (**Li et al., 2016**). Among those PRR, MDA5 (Melanoma Differentiation-Associated protein 5) and RIG1 (Retionic Acid Inducible Gene I) are one of the main protein sensors of the pathogen-associated molecular patterns (PAMPs), functioning as sensors of RNA viruses, and help to identify some DNA viruses **(Kell and Gale Jr, 2015),** in the form of viral double-stranded RNA (dsRNA) motifs to induce expression of type 1 interferons (IFN1) (IFNα and IFNβ) and other pro-inflammatory cytokines during the early stage of viral infection (**Brisse1 and Ly, 2019**). When viral nucleic acid enters into the endosome firstly get countered by TLRs potently which stimulates innate immune responses through the rapid action of IFNα/β-dependent and independent pathways (**Thompson and Iwasaki, 2008**) as a response of expression of duck RIG-I gene (Huo et al. 2019). We had previously sequenced certain immune response genes as TLR2, TLR3, TLR7 and RIGI from ileocaecal junction or tonsil of Bengal duck (Pal et al., 2017a,b,c,d)

Duck has varied eco-genetic potentials which could undoubtedly bring the up-liftment of rural livelihoods in the country. Duck eggs and meat are accepted by all sections of the society. About 96.47% of ducks are reared in rural areas while the remaining 3.53% are reared in urban areas. Since majority of ducks are desi or non-descript type having low production potential, therefore, there is an ample scope of duck improvement in the region.

The study upon indigenous type of ducks ( **Kang et al., 2014, Pal et al., 2020)** stated that they are more resistant to many diseases such as Newcastle disease, than that of poultry, despite their frequent exposure to marshy and grazing areas where the incidence of potential pathogens is relatively high and this might be due to the strong innate immune response. In spite of comparatively better resistant to common poultry disease certain diseases can infect duck like duck plague. The disease is characterized by certain death, high mortality, hemorrhage and necrosis in internal organs.

Replication of duck plague virus is fast and cell-to-cell spread of the infectious agent that is the virion, does not require the development of envelop (**Yu You, 2018**). Quantitative real-time PCR demonstrate the innate immunity related genes may show the positive results in various organs. Although duck plague is an important economic disease of duck, the studies on innate immunity of duck with respect to duck plague at molecular level is scares except a single study by **Li et al.(2016)** where they have studied an brain and spleen. In the current study we proposed to study the molecular pathogenesis of different body organs including gut and other organs which are affected by duck plague virus. In our lab, we had already studied the effect of immune response genes against avian influenza, whose genome content is RNA (Pal et al., 2021). Here we attempt to study immune responsiveness against DNA virus as Duck plague virus.

Hence, we aim to explore the gut associated lymphoid tissue and to explore the role of important immune response genes as RIG1, MDA5 and interferon alpha genes in important organs such as liver in both healthy and duck plague infected duck in both in silico approach followed by experimental validation.

## Materials and Methods

In the present study, we have studied the role of three important host immune response genes (RIGI, MDA5 and Interferon alpha) of duck and their role in mediating duck plague viral infection. RIGI and MDA5 are the genes for receptor molecules, which in turn releases interferon alpha, thus acting against duck plague viral infection.

In the first step, we characterize these genes of Bengal duck through cloning, sequencing and *in silico* studies to identify the important domains. In the next step, we performed molecular docking analysis for four important proteins of Duck plague virus of West Bengal origin glycoprotein (_MH745151.1_), UL30 (_LC105645_), UL31(_KJ451479_), thymidine kinase (_KF214787.1_) with the host immune response gene (RIGI, MDA5 and Interferon alpha). Glycoprotein being the structural protein of duck plague virus, very high patchdock score was observed with RIGI, MDA5, which are the receptor molecules.

Challenge studies with duck plague viral infection to Bengal ducks were carried out in two groups-challenged and control. Infected subjects were confirmed through molecular detection and viral load was estimated. Differential mRNA expression profiling was studied for confirmation of the findings in infected and control samples. Differential mRNA expression profiling for these genes specific to host resistance were carried out with respect to different GALT tissues. Significantly better expression was observed in liver; hence futher studies were conducted with liver tissue for comparison of infeted vs. healthy duck samples. Infected ducks were detected through symptomatic diagnosis, PCR based molecular diagnosis. Haematological and biochemical parameters were carried out for both infected and healthy groups. Histological section of liver and kupfer cells were studied followed by immunohistochemistry with respect to these proteins.

### Characterization of RIGI, MDA5 and interferon alpha gene in duck

#### Birds and sample collection

The samples were collected from Bengal ducks maintained at West Bengal University of Animal and Fishery Sciences. Tissue samples were also collected from slaughter house for Bengal duck.

#### Sample Collection and RNA Isolation

Duck liver tissue (1g) was collected from slaughtered birds which were clinically healthy and maintained in duck house of LFC Dept, WBUAFS. Adult males (n=6) were selected for collection of samples. Tissue was immersed in Trizol in vial and transported in ice to the laboratory for RNA isolation. Total RNA was isolated using TRIzol extraction method (Life Technologies, USA), as per the standard procedure and was further utilized for cDNA synthesis (**Pal and Chatterjee, 2009; Pal et al., 2011, Banerjee et al., 2021, Rawat et al., 2021).** cDNA concentration was estimated and samples above 1200 microgram per ml were considered for further study.Tissues from gut associated lymphoid tissue (GALT) as ileocaecal junction, liver, gizzard, caecum, jejunum, spleen, pancreas were collected from both healthy and duck infected with duck plague virus for studying the expression profiling with quantitative PCR.

#### cDNA synthesis and PCR Amplification of the RIGI, MDA5 and interferon Gene of Bengal Duck

20μL of reaction mixture was composed of 5μg of total RNA, 0.5μg of oligo dT primer (16–18mer), 40U of Ribonuclease inhibitor, 1000M of dNTP mix, 10mM of DTT, and 5U of MuMLV reverse transcriptase in appropriate buffer. The reaction mixture was mixed thoroughly followed by incubation at 37°C for 1 hour. The reaction was allowed upto 10 minutes by heating the mixture at 70°C unliganded and then chilled on ice. Afterwards, the integrity of the cDNA was checked by performing PCR (Pal, A, 2021, Pal and Chatterjee,2010, Pal et al., 2014a,b, Pal et al., 2019ab, Pal et al., 2020, Pal et 2011). Concentration of cDNA was estimated through Nanodrop. RIGI, MDA5 and interferon gene primer pairs were designed based on the published sequences of chicken and duck origin using DNASTAR software (Hitachi Miraibio Inc., USA) to amplify full length open reading frame (ORF) of these genes (Table 1).

**Table 1:**
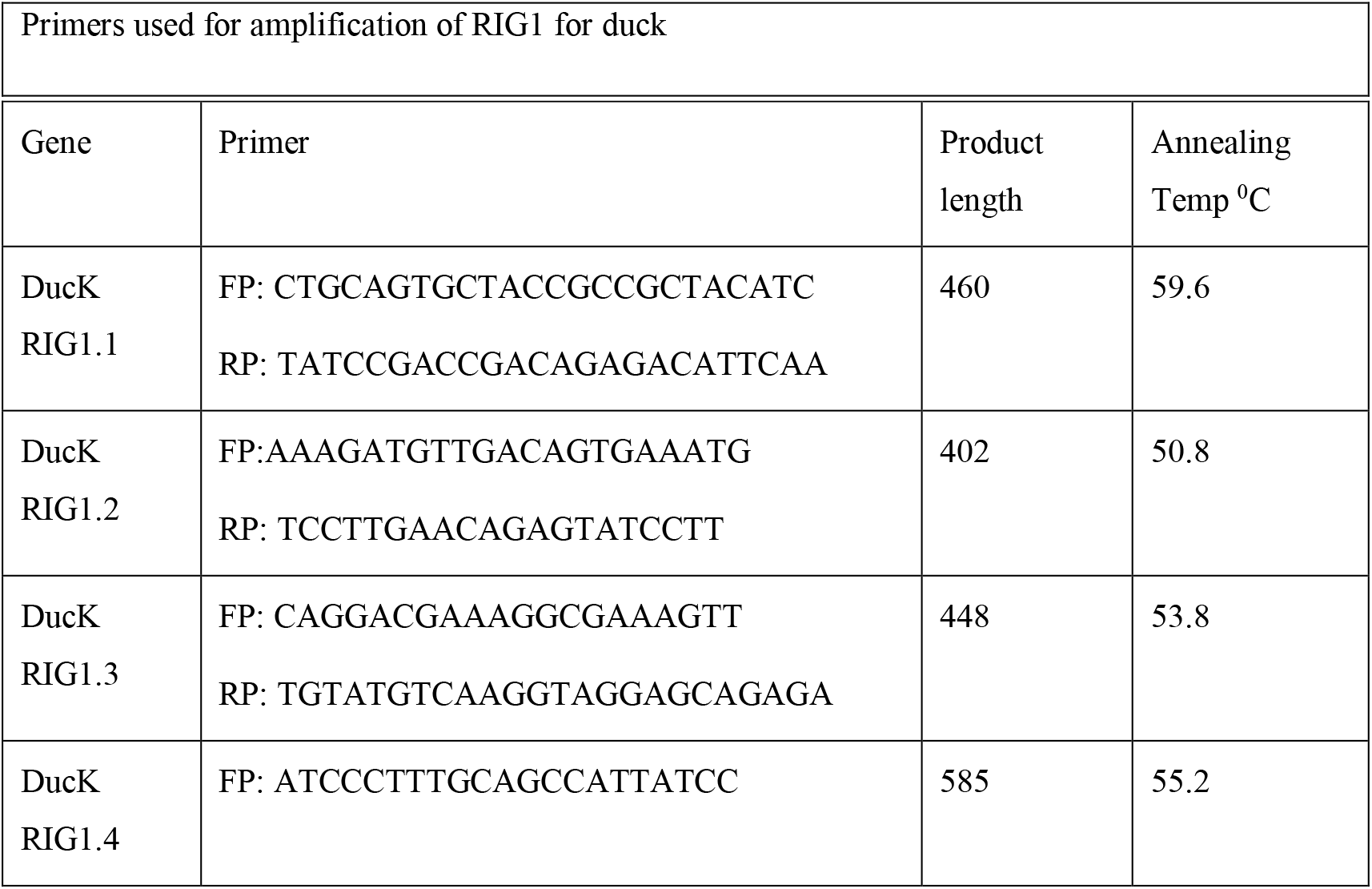
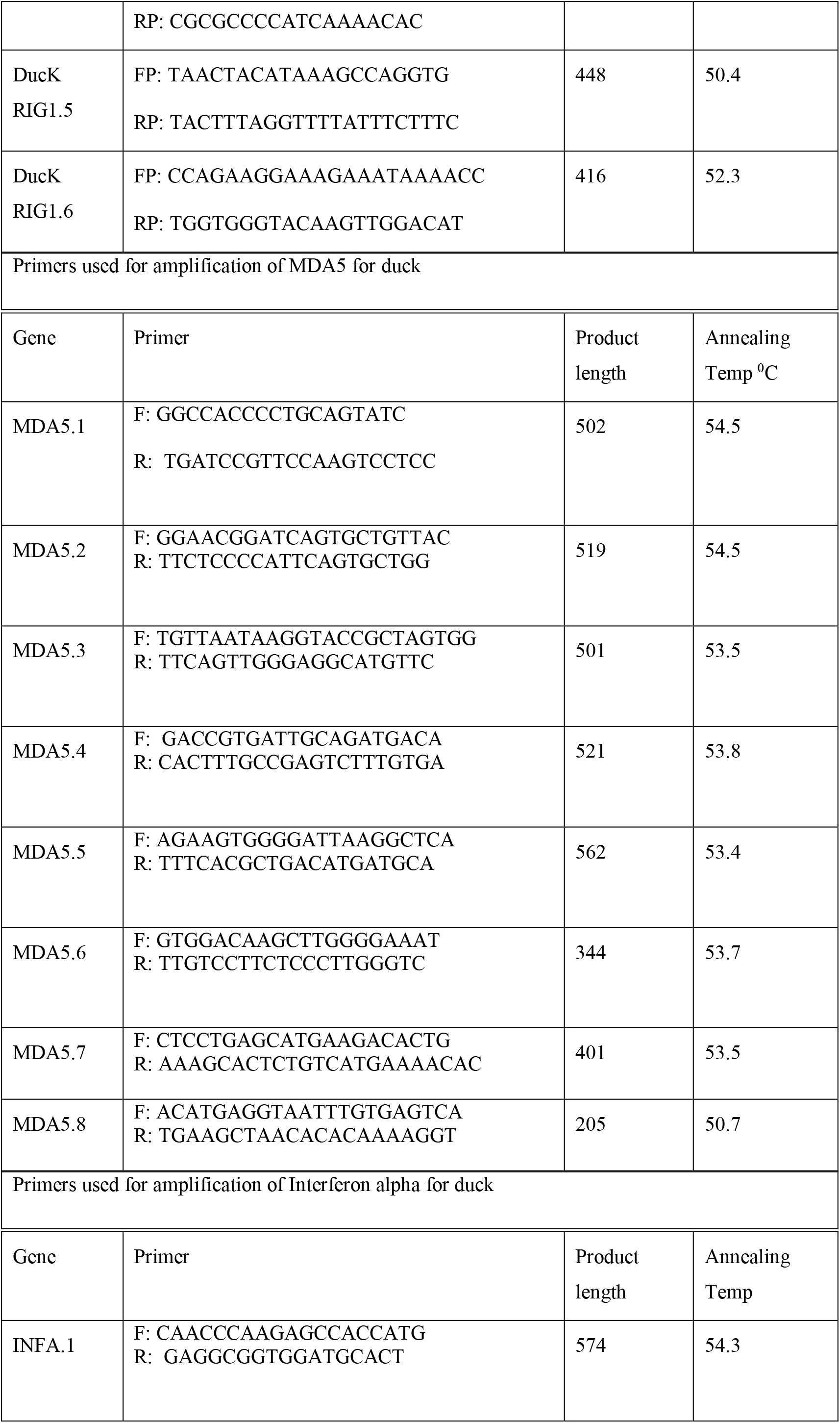
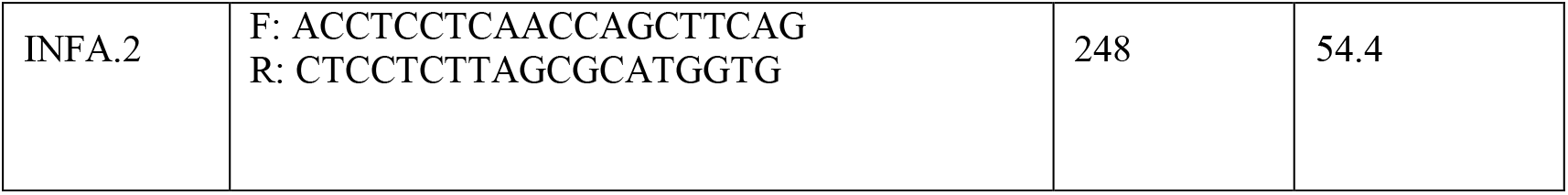
List of primers used for the amplification of immune response genes.

25μL of reaction mixture was comprised of 80–100ng cDNA, 3.0μL 10X PCR assay buffer, 0.5μL of 10mM dNTP, 1U Taq DNA polymerase, 60ng of each primer, and 2mM MgCl2. PCR-reactions were performed in a thermocycler (PTC-200, MJ Research, USA) with cycling conditions as, initial denaturation at 94°C for 3min, further denaturation at 94°C for 30sec, annealing at 61°C for 35sec, and extension at 72°C for 3min were conducted for 35 cycles followed by final extension at 72°C for 10 min.

#### cDNA Cloning and Sequencing

Amplified products of RIGI, MDA5 and interferon alpha were checked by 1% agarose gel electrophoresis. The products were purified from gel using Gel extraction kit (Qiagen GmbH, Hilden, Germany) .pGEM-T easy cloning vector (Promega, Madison, WI, USA) was used for cloning. Then, 10μL of ligated product was mixed thoroughly to 200μL competent cells, and heat shock was given at 42°C for 45sec in a water bath. Subsequently, the cells were immediately transferred on chilled ice for 5 min., and SOC medium was added to it. The bacterial culture was centrifuged to obtain the pellet and plated on LB agar plate containing Ampicillin (100mg/mL) added to agar plate @1:1000, IPTG (200mg/mL) and X-Gal (20mg/mL) for blue-white screening. Plasmid isolation from overnight-grown culture was carried out by small-scale alkaline lysis method as described in (Sambrook et al., 2001). Recombinant plasmids were characterized by PCR using CD14 primers as reported earlier and restriction enzyme digestion. CD14 gene fragments released by enzyme EcoRI (MBI Fermentas, USA) were inserted in recombinant plasmid which was sequenced by dideoxy chain termination method with T7 and SP6 primers in an automated sequencer (ABI prism, Chromous Biotech, Bangalore).

#### Sequence Analysis

DNASTAR Version 4.0, Inc., USA software was employed for the nucleotide sequence analysis for protein translation, sequence alignments, and contigs comparisons (Pal et al.,2004,Pal et al., 2005). Novel sequences was submitted to the NCBI Genbank and accession numbers were obtained which is available in public domain.

#### Study of Predicted duck RIGI, MDA5, IFN alpha Protein Using Bioinformatics Tools

Predicted peptide sequences of duck RIGI, MDA5, IFN alpha Protein were then aligned with that of other species using MAFFT (**Katoh et al.,, 2013**). The analysis was conducted for a sequence based comparative study. Signal peptide is essential to prompt a cell to translocate the protein, usually to the cellular membrane and ultimately signal peptide is cleaved to give mature protein. Prediction of presence and location of signal peptide of these gene were conducted using the software (SignalP 3.0 Sewer-prediction results, Technical University of Denmark). Leucine percentage was calculated manually from predicted peptide sequence. Di-sulphide bonds are essential for protein folding and stability, ultimately. It is the 3D structure of protein which is biological active.

Protein sequence level analysis was employed (http://www.expasy.org./tools/blast/) for assessment of leucine rich repeats (LRR), leucine zipper, detection of Leucine-rich nuclear export signals (NES), and detection of the position of GPI anchor, N-linked glycosylation sites. Since these are receptors, these are rich in leucine rich repeat, which are essential for pathogen recognition and binding. Leucine zipper is essential to assess dimerization of IR molecules. N-linked glycosylation is important for the molecule to determine its membranous or soluble form. Leucine rich nuclear export signal is essential for export of these protein from nucleus to cytoplasm, whereas GPI anchor is responsible for anchoring in case of membranous protein. Prosite was used for LRR site detection.

Leucine-rich nuclear export signals (NES) was analyzed with NetNES 1.1 Server, Technical university of Denmark. O-linked glycosylation sites were detected using NetOGlyc 3.1 server (http://www.expassy.org/), whereas N-linked glycosylation site were assessed through NetNGlyc 1.0 software (http://www.expassy.org/). Sites for leucine-zipper were detected through Expassy software, Technical university of Denmark (**Glick, 1977**). Sites for alpha helix and beta sheet were detected using NetSurfP-Protein Surface Accessibility and Secondary Structure Predictions, Technical University of Denmark (**Petersen et al., 2009).** Domain linker site were predicted (**Ebina et al., 2009**). LPS-binding (**Cunningham et al., 2000**) and LPS-signalling sites **(Muroi et al., 2002)** were predicted based on homology studies with other species CD14 polypeptide. These sites are important for pathogen recognition and binding.

#### Three dimensional structure prediction and Model quality assessment

Three dimensional model of RIGI, MDA5 and Interferon alpha polypeptide were predicted through Swissmodel repository (**Kiefer et al., 2009**). Templates possessing greatest identity of sequences with our target template were identified with PSI-BLAST (http://blast.ncbi.nlm.nih.gov/Blast). PHYRE2 server based on ‘Homology modelling approach’ was used to build three dimensional model of these proteins (**Kelley, 2015**). Molecular visualization tool as PyMOL (http://www.pymol.org/) was employed for model generation and visualization of three dimensional structure of these protein understudy for duck origin. The structure of duck molecules were evaluated and assessed for its stereochemical quality (through SAVES, Structural Analysis and Verification Server, http://nihserver.mbi.ucla.edu/SAVES/); then refined and validated through ProSA (Protein Structure Analysis) web server (https://prosa.services.came.sbg.ac.at/prosa) (**Wiederstein and Sippl, 2007**). NetSurfP server (http://www.cbs.dtu.dk/services/NetSurfP/ **Peterson et al., 2009)** was used for assessing surface area of these protein through relative surface accessibility, Z-fit score, and probability of alpha-Helix, beta-strand and coil score.

Alignment of 3-D structure of these proteins were analyzed with RMSD estimation to evaluate the structural differentiation by TM-align software (**Zhang et al., 2005**).

#### Protein-protein interaction network depiction

In order to understand protein interaction network of these proteins, we performed search in STRING 9.1 database (**<u>Franceschini</u> et al., 2015**). The functional interaction was assessed with confidence score. Interactions with score < 0.3, scores ranging from 0.3 to 0.7, and scores >0.7 are classified as low, medium and high confidence respectively. Also, we executed KEGG analysis which depicted the functional association of these proteins with other related proteins.

### *In silico* study for the detection of binding site of RIGI, MDA5, Interferon alpha with *Duck plague virus*

#### Molecular Docking of host RIGI, MDA5 and Interferon alpha with the viral proteins for molecular diagnosis

Molecular docking is a bioinformatics tool used for *in silico* analysis for the prediction of binding mode of a ligand with a protein 3D structure. Patch dock is an algorithm for molecular docking based on shape complementarity principle (Duhovny **et al., 2005**). Patch Dock algorithm was used to predict ligand protein docking for surface antigen for four important proteins of Duck plague virus of West Bengal origin (glycoprotein, UL30, UL31, thymidine kinase) with the host immune response gene (RIGI, MDA5 and Interferon alpha). Firedock analysis was assessed for further confirmation.

#### Challenge study with DP virus, symptomatic diagnosis, estimation of infective viral load at different stages of infection, PCR based detection

For the purpose of challenge study, initially we had incubated the fertile duck eggs (n=12) in incubator at 37°C along with water in a petridish in order to maintain the proper humidity, which is essential for incubation of duck eggs. After one week, we rechecked and screened the fertile embryo through candling. In order to harvest the duck plague virus, at nine days of incubation of the fertilized egg, we puncture the air sac with the help of a sterilized needle through the chorioallantoic membrane, and innoculated 200ul of the infective agent in a sterile environment. The needle puncture was quickly sealed with feviquick. The eggs were again incubated in sterile environment 37C and humid condition as earlier. The entire procedures were undertaken at BSL3 lab. Duck plague viral strain employed was the available infection in the state from the clinical infected cases. After 3 days of innoculation, the viruses was isolated from embryonic fibroblast cells through viral DNA isolation kit. Quantification of viral load was estimated through quantitative PCR with duck plague vaccine as control. For the purpose of challenging, 0.3ml of viral stock at EID50 (Egg infective dose 50) was employed to each bird I/M. Infective amnio-allantoic fluid was used to determine 50 percent EID. Initial verification was conducted to assure that the ducks were free from DP viral infection.

#### Sample Collection

For the present study six tissue samples were collected from the infected indigenous duck samples followed by challenge study. We designate the ducks with mortality as infected or diseased (less immuned) and the ducks that survived as healthy (better immuned).

#### Diagnosis and confirmation for duck plague

The ducks were initially diagnosed as duck plague clinically with the help of initial symptoms. Later on confirmatory diagnosis was conducted through molecular detection of Duck plague virus in the infected samples.

#### Gross Anatomical view of the organs in duck plague infected duck

We studied different body organs from infected duck for clinical diagnosis of duck plague.

#### Confirmation through molecular detection

In the next step, we follow confirmation of the samples with molecular PCR based technique using the following primers as recommended by OIE, Paris.

It includes viral DNA isolation through isolation kit and subjected to PCR. We isolate DNA since Herpes virus is a DNA virus.

#### Assessment of dose of viral particles

We estimated the dose of viral particles standardization through Quantitative PCR. We take the standard duck plague vaccine as a control for its amplification and quantification.

**Challenge study with Duck Plague virus and study of expression profiling in healthy and infective groups.**

**Differential mRNA expression profiling for GALT tissues w.r.t. RIGI, MDA5 and Interferon alpha gene.**

**RT-PCR (Real Time-Polymerase Chain Reaction)**

#### RNA Estimation

The tissue samples were cut into small pieces and submerged in liquid Trizol followed by thorough grinding with a mortar and pestle. 1 ml triturated tissue was taken into 2 ml eppendorf tube and 1 ml chloroform was added. Then it was centrifuged at the rate of 10,000 rpm for 10 minutes at 4°C in automated temperature controlled refrigerated centrifuge machine (BR Biochem, Life Sciences and REMI C24 plus). Three phases differentiation were identified. Then upper most aqueous phase was collected for RNA isolation into a new eppendorf tube and equal volume of 100% isopropanol was added. It was left for a minute at 20-25°C temperature and centrifuged at the rate of 10,000 rpm for 10 minutes. After that supernatant was discarded and equal volume of 70% ethanol was added to the pellet. Again the mixture was centrifuged at the rate of 10,000 rpm for 10 minutes and then supernatant was discarded. Then the eppendorf tube containing pellet kept at room temperature for air drying. After drying the remaining pellet dissolved with nuclease free water (Applied Biosystems, Cat. No. AM9930) and stored at −20°C for future uses **(Pal et al., 2011, Pal, 2009, Pal et al., 2014).**

#### Qualitative Analysis of Total RNA

The total RNA was quantified using NanoDrop-8000 (Thermo Scientific, Model No. 8000 Spectrophotometer), taking 1μl of each sample to determine the concentration and A260/280 ratio (Rawat et al. 2021, Banerjee et al., 2021, Pradhan et al., 2018).

#### First Strand cDNA Synthesis (Reverse Transcriptase -PCR)

5μl estimated RNA was taken into a PCR micro tube and added 1 μl OligoDT primer, 1μl 10mM dNTP and 13μl distilled water. After well mixing heated at 65°C for 5 minutes. Then those tubes were quickly chilled on ice for 5-7 minutes and centrifuged. Then 4 μl 5x first strand buffer and 2 μl DTT (100mM) were added to the tube. It was well mixed and incubated at 37°C for 2 minutes. Then 1 μl M-MLVRT (200 u/μl) added to the reaction mixture and treated at 37°C for 50 minutes and at 70°C for 15 minutes for inactivation. All the temperature treatments were maintained automatically into the thermal cycler machine (Applied Bio systems by Life Technologies) by setting the whole programme.

#### Primer Designing and Synthesis

All the primers were designed using primer 3 software (v. 0. 4.0) as per the recommended criteria. MDA5, RIG1 and GAPDH used in the experiment, were designed (DNASTAR software).

#### Real Time PCR Experiment (SYBR Green based)

The primers were standardising with respective cDNA sample before subjected to real time PCR. The entire reactions were performed in triplicate (as per MIQE Guidelines) and the total volume of reaction mixture was set up to 20μl. The reaction mixture is set up with MDA5, RIG1 and GAPDH (housekeeping gene) primers calculated as per concentration. 1μl of cDNA, 10μl Hi-SYBr Master Mix (HIMEDIA MBT074) and rest volume adjusted with nuclease free water to achieve the total reaction mixture volume. Then the reaction plate placed into the ABI 7500 system and run the reaction program. The delta-delta-Ct (ΔΔCt) method used for analysis the result. The primers used for reaction are followed:

### Estimation of haematological and biochemical parameter

#### Haematological Profiles

The haematological parameters like hemoglobin, erythrocyte sedimentation rate (ESR) and packed cell volume (PCV) were estimated in whole blood soon after the collection of blood. Hemoglobin was estimated by acid haematin method (**Benjamin, 1985**), E.S.R. and PCV by Wintrobe’s tube (**Hawk, 1965**). The total erythrocyte count (TEC), total leukocyte count (TLC) and Differential leukocyte count (DLC) were studied by standard methods described by **Jain (1986).**

#### Biochemical Analysis

The serum biochemical parameters, estimated in the experiments were total protein, albumin, globulin, albumin: globulin, aspartate aminotransferase (AST), alanine aminotransferase (ALT), alkaline phosphatase (ALP), Total bilirubin, Indirect bilirubin, direct bilirubin, glucose, uric acid, urea and BUN by using a semi-auto biochemistry analyzer (Span diagnostic Ltd.) with standard kits (Trans Asia Bio-Medicals Ltd., Solan, HP, India). The methodology used for estimation of total protein, albumin, total & direct bilirubin, ALT, ALP, glucose, creatinine urea and uric acid were biuret method, bromocresol green (BCG) method, 2-4-DNPH method, modified kind and king’s method, GOD/POD method, modified Jaffe’s Kinetic method. GLDH-urease method and trinder peroxidise method respectively.

#### Histological section

The liver samples were fixed in formalin (10%) and embedded in paraffin and processed for histological examination and stereology. The liver tissues were submerged in Lillie fixative for 1 week at room temperature and then were processed and embedded vertically in paraffin wax. Then, each liver sample was exhaustively sectioned into 4 μm-thick sections by a fully automated rotary microtome (Leica RM2255, Germany). Each of these sections was stained with hematoxylin and eosin and mounted. From each liver sample, 10-15 sections were chosen by systematic uniform random sampling (SURS) method.

#### Immunohistochemistry

Paraffin tissue blocks were prepared by standard manual alcohol-acetone protocol. The 5-6 μm thick paraffin sections obtained from different segments of the uterus (middlehorn of uterus and tip of the horn) of both the pregnant and non-pregnant ewes, were taken on Millennia 2.0 adhesion slides (Cat. No. 71863-01, abcam).The tissue sections were deparaffinized and hydrated in distilled water. The tissues slides were covered with trypsin enzymatic antigen retrieval solution (Cat. No. ab970, abcam) and kept in incubator in humid environment at 37°C for 5-10 minutes. The sections were then incubated for 60 minutes in peroxidase blocking solution (Lot. No. 00065614, Dako) at room temperature to block non-specific antibody binding activity. After subsequent washing with phosphate buffer saline (PBS), the sections were incubated at 37°C for 2 hours in humid environment with mouse monoclonal anti RIGI antibody and MDA5 antibody in 1: 200 dilution. Immunoreactivity was detected after one hour incubation at 37°C with secondary antibody, Rabbit anti-mouse IgG H&L (HRB Conjugated, Cat. No. ab6728; abcam) in dilution 1:200. Slides were then rinsed 3 times in PBS for 5 minutes each, followed by treatment with freshly prepared DAB solution for 3 minutes (DAB substrate, Cat. No. 34001, Thermo Fisher Scientific). The sections were counter stained with Mayer’s haematoxylin, hydrated in ethanol, cleared in xylene and then mounted in DPX.

#### Statistical Analysis

Microsoft Excel used for the descriptive statistical analysis. SYSTAT 13.1 software (SYSTAT Software Inc.) used for statistical analysis and analysis of variance (ANOVA) was used to test between parameters.

## Results

### Characterization of Duck RIGI, MDA 5, Interferon alpha

First at molecular level, we had characterized the genes and assessed the differential mRNA expression profiling of certain genes as RIGI, MDA5, Interferon alpha pertaining to tissues from different body organs. Sequences are submitted to NCBI. Published sequences are RIGI (Acc no.**OL438907**). 3D structure for RIGI, MDA5 and Interferon alpha for duck have been predicted in Fig 1, Fig 2 and Fig 3 respectively. RIGI is an important gene conferring antiviral immunity. A series of post-translational modification and various domains for its important function have been represented.

**Fig 1:**
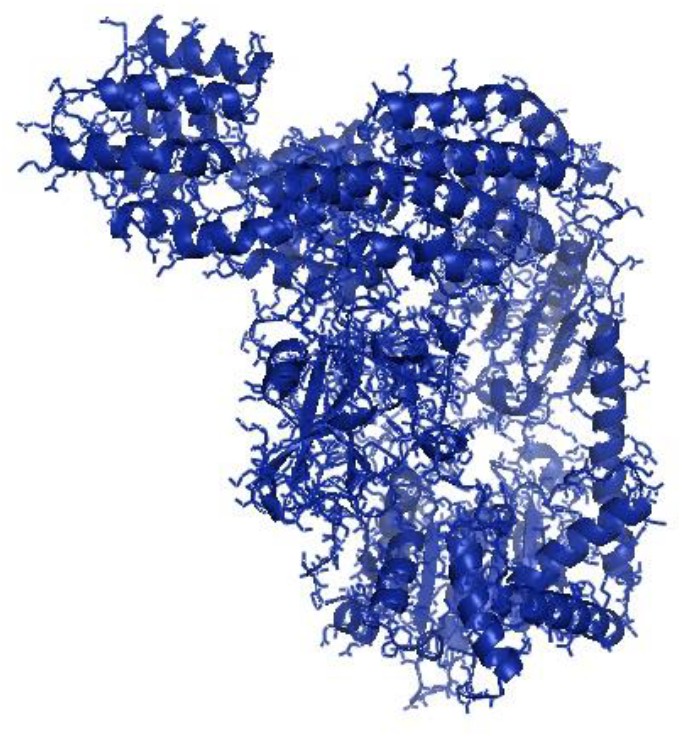
3D structure of duck RIGI molecule.

**Fig 2:**
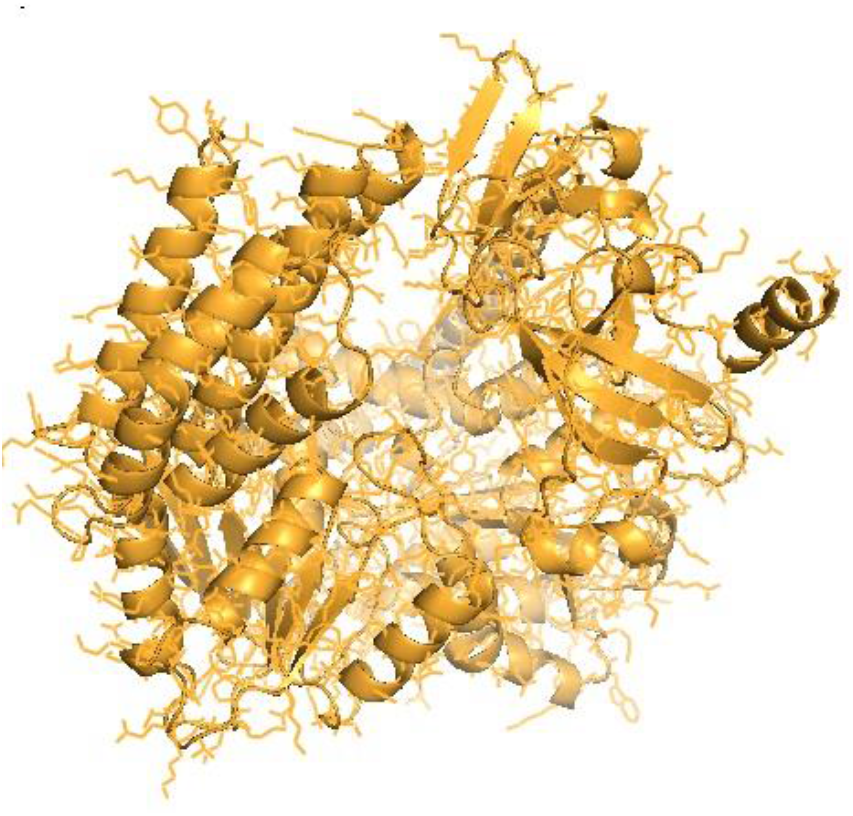
3D structure of duck MDA5 molecule.

**Fig 3:**
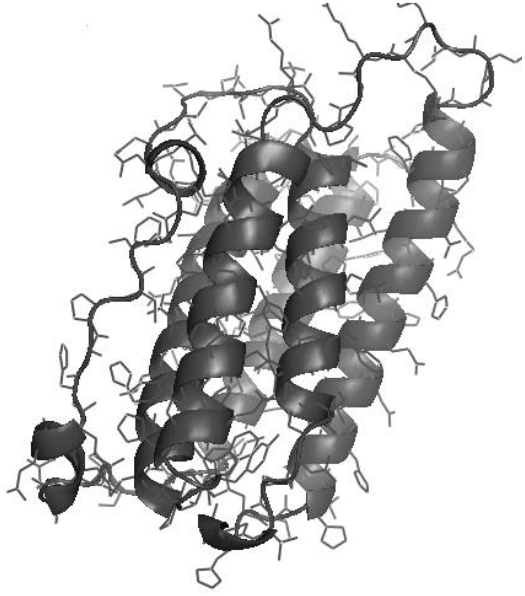
3D structure of duck interferon alpha molecule.

CARD_RIG1 (Caspase activation and recruitment domain found in RIG1) have been depicted in amino acid position 2-91(hot pink), 99-188(fire brick red) Fig 4a. CARD2 interaction site (17-20 orange, 23-24 green, 49-50 chocolate, 79-84 yellow) Fig 4b, CARD1 interface (100 red, 103 warm pink, 130-135 yellow orange, 155 ruby, 159 magenta, 161-162 limon) Fig 4b, helical insert domain interface (101 red, 104-105 green, 107-108 yellow, 110-112 orange, 114-115 hot pink, 139 chocolate, 143-145cyan, 147-148grey, 151 magenta, 180 firebrick, 183-184 forest, 186 aa yellow orange) have been depicted at RIG1 of duck Fig 4c. Fig 4d depicts helicase insert domain (242-800 aa) as yellow, helicase domain interface (polypeptide binding) as (511-512aa red, 515aa green, 519aa orange). Fig 4e depicts double-stranded RNA binding site (nucleotide binding) at amino acid positions 832 (red), 855 (green), 876-877 (yellow), 889-891(magenta), 911 (white). The sites for RD interface (polypeptide binding) and RIGI-C (C terminal domain of retinoic acid-inducible gene, RIG-I protein, a cytoplasmic viral RNA receptor) have been depicted in Fig 4f. The site for RIG-I-C as amino acid position 807-921 represented by a mesh of green tints. The sites for RD interface have been depicted as amino acid positions 519 (red sphere), 522-523 (magenta), 536-537 (orange), 540 (green). The sites for RNA binding have been depicted as 511-512 (hot pink), 515 (yellow orange), 519 (green) in Fig 4h.

**Fig 4a:**
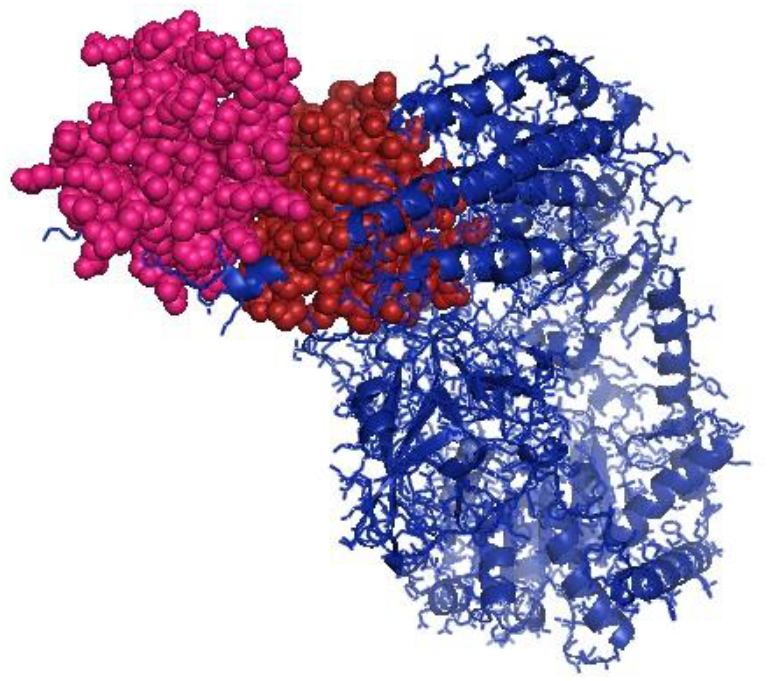
CARD_RIG1 (Caspase activation and recruitment domain found in RIG1) have been depicted in amino acid position 2-91, 99-188.

**Fig 4b:**
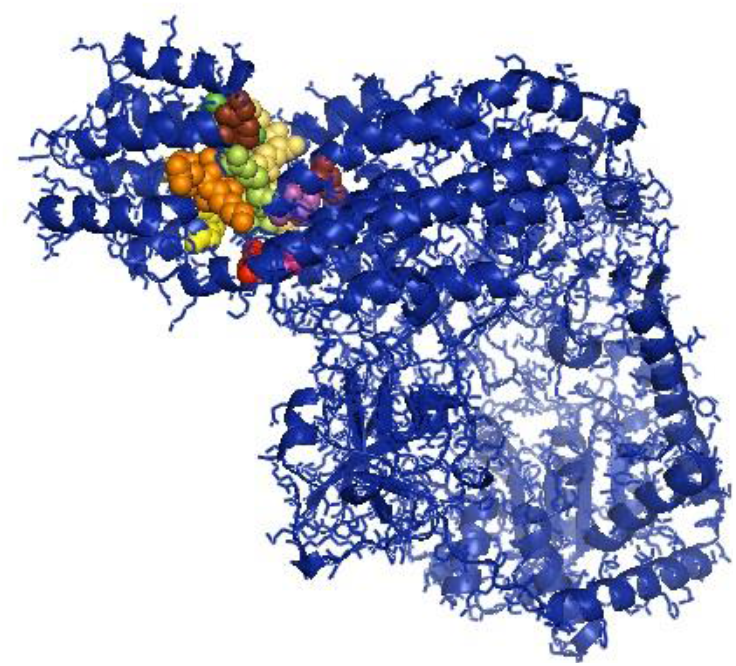
CARD2 interaction site (17-20, 23-24, 49-50, 79-84), CARD1 interface (100, 103, 130-135, 155, 159,161-162)

**Fig 4c:**
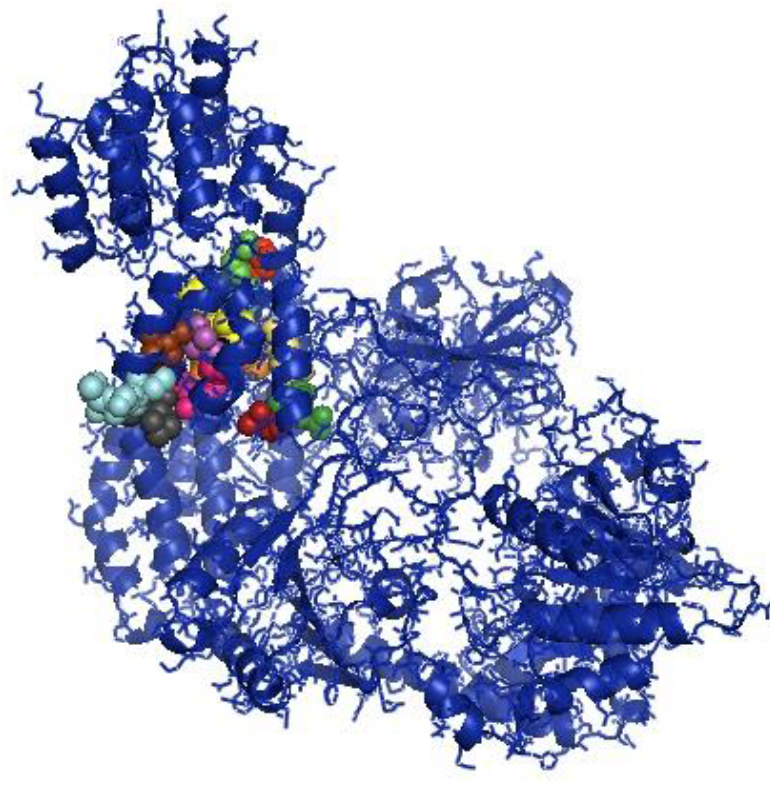
Helical insert domain interface (101, 104-105, 107-108, 110-112, 114-115,139, 143-145, 147-148, 151, 180, 183-184, 186 aa) have been depicted at RIG1 of duck.

**Fig 4d:**
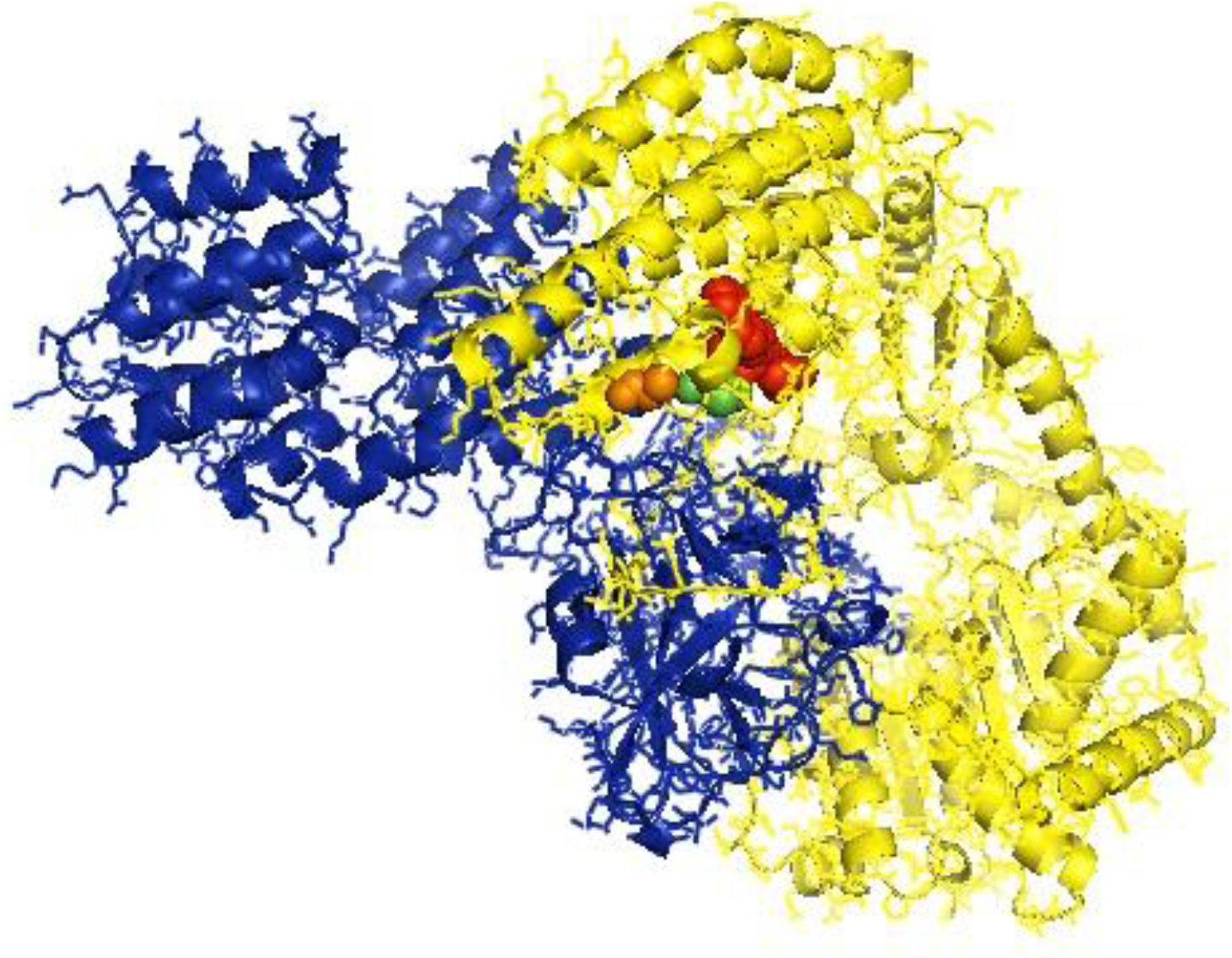
Helicase insert domain (242-800 aa), helicase domain interface (polypeptide binding) as (511-512, 515, 519 aa).

**Fig 4e:**
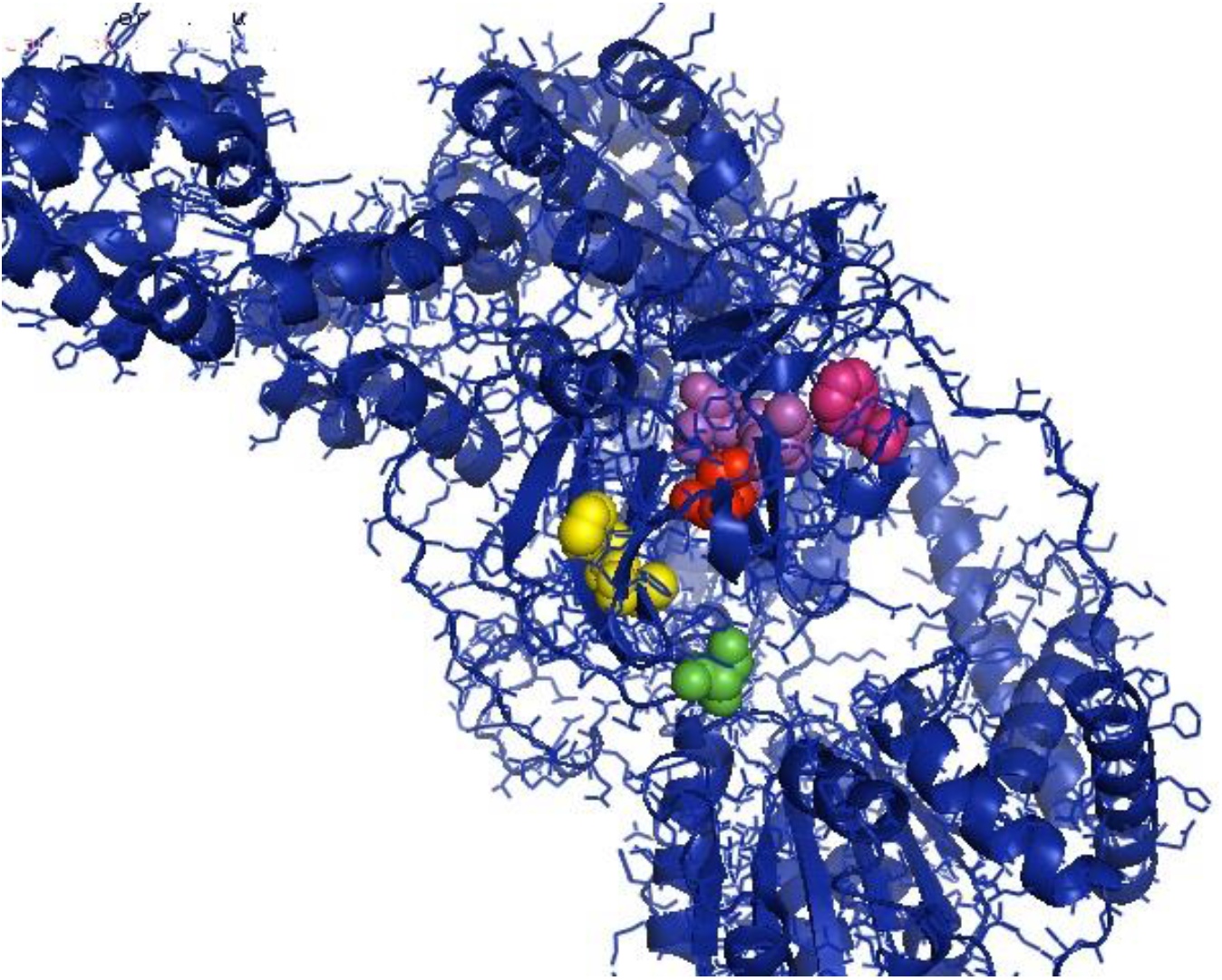
Double-stranded RNA binding site (nucleotide binding) at amino acid positions 832, 855, 876-877, 889-891, 911.

**Fig 4f:**
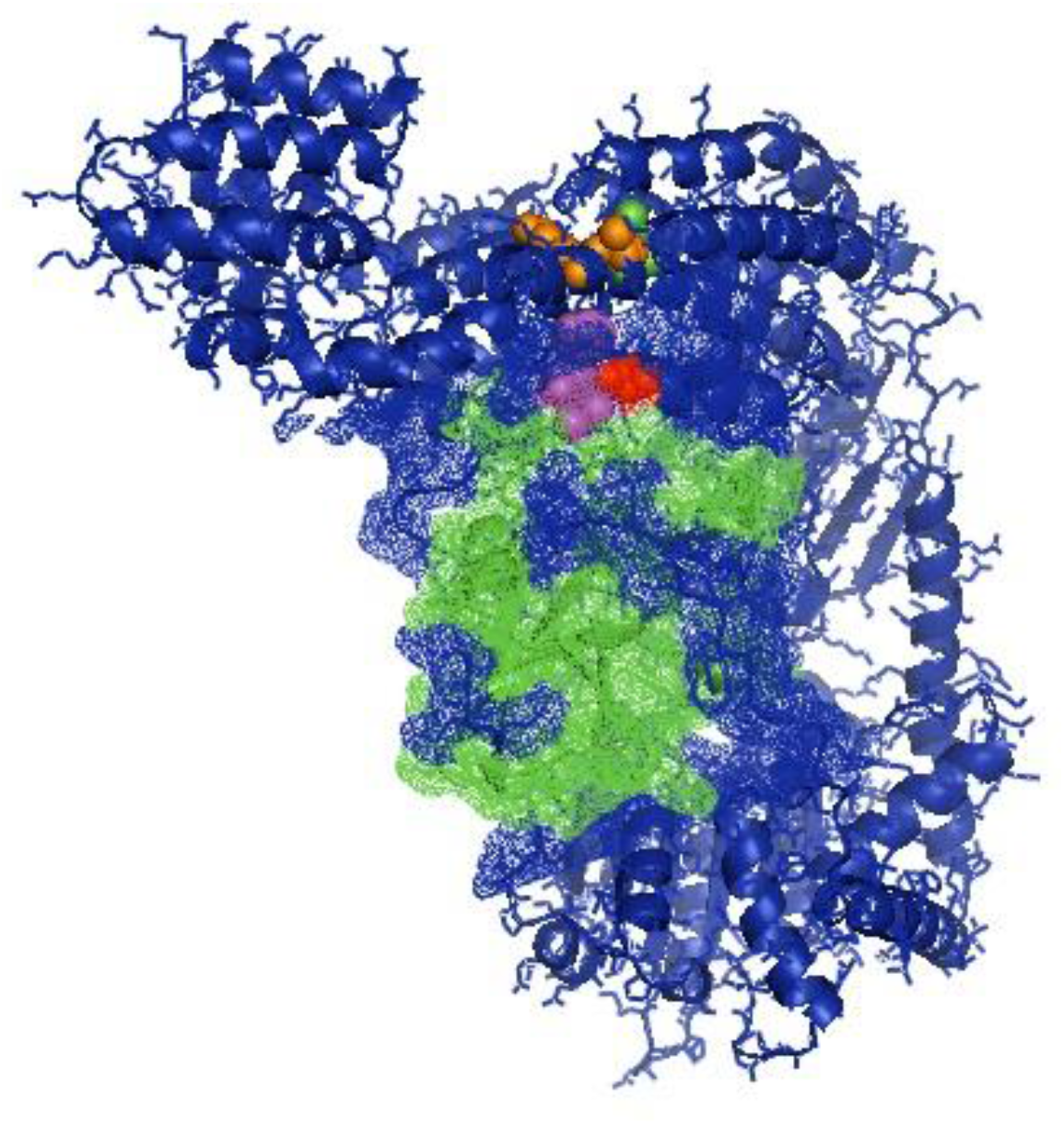
The sites for RD interface (polypeptide binding) and RIGI-C (C terminal domain of retinoic acid-inducible gene, RIG-I protein, a cytoplasmic viral RNA receptor)

**Fig 4g:**
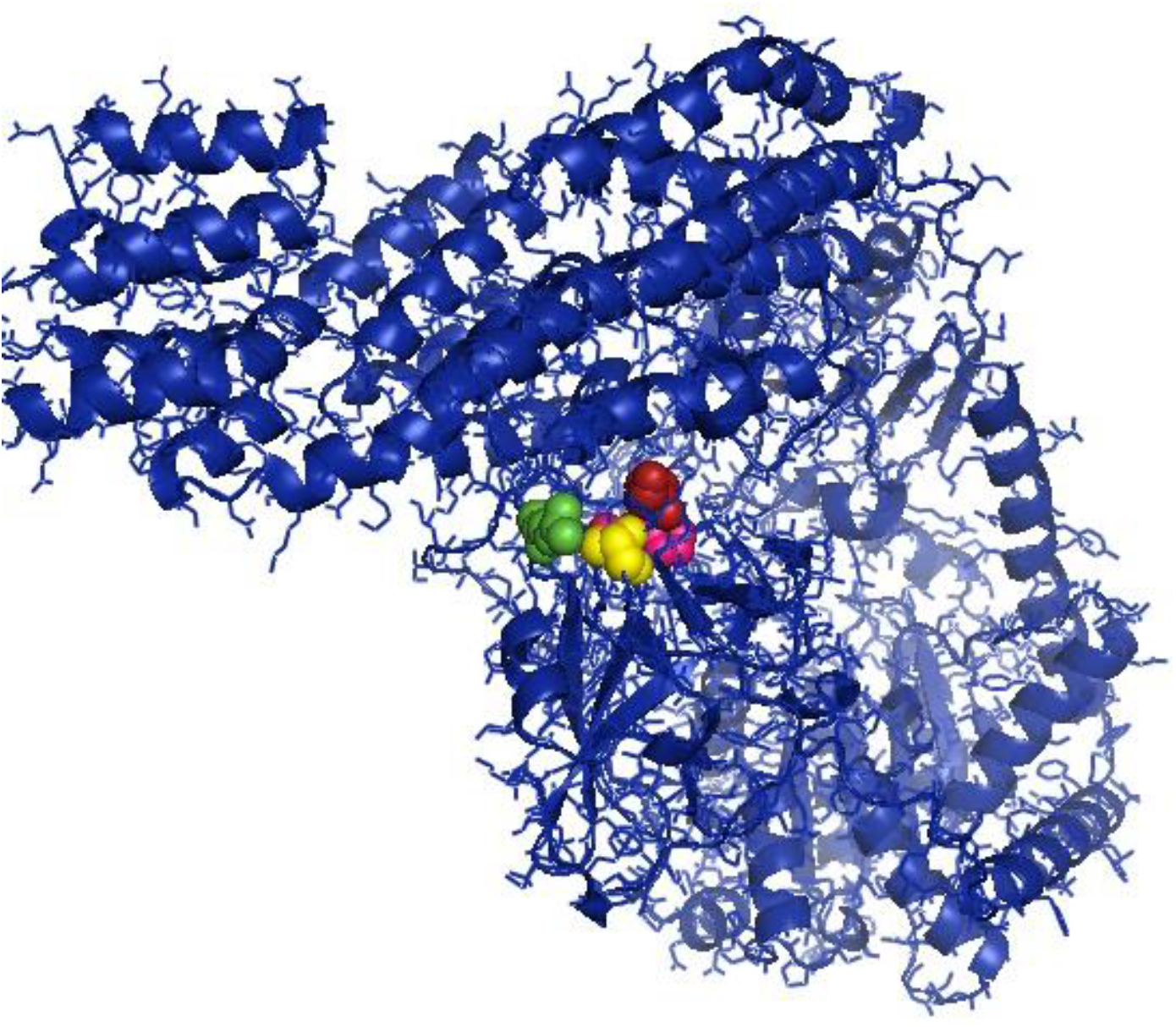
Site for RIG-I-C as amino acid position 807-921. The sites for RD interface at amino acid positions 519, 522-523, 536-537, 540.

**Fig 4h:**
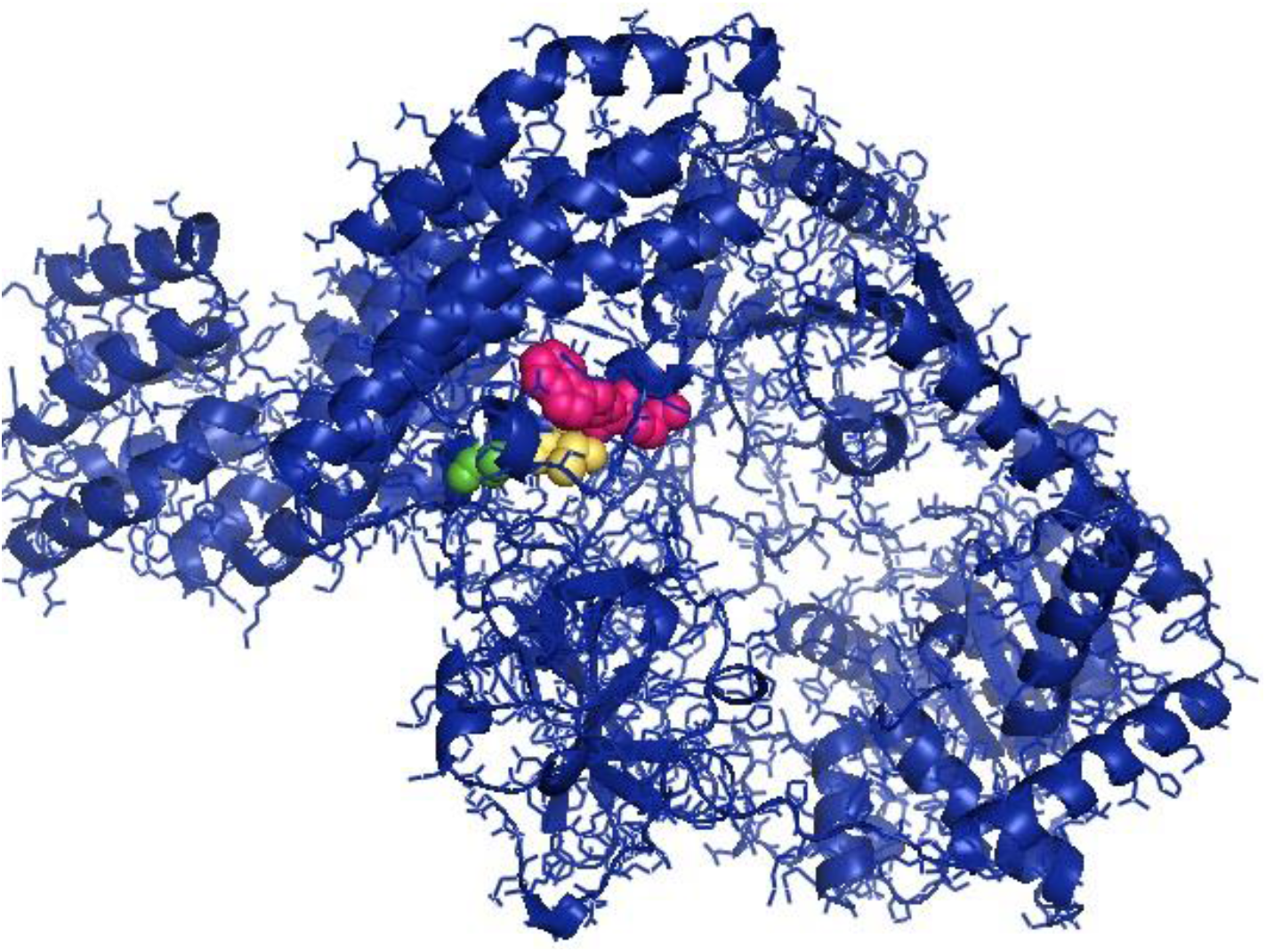
The sites for RNA binding have been depicted as 511-512, 515, 519.

Certain important domains have been identified in duck MDA5. Helicase ATP binding site was predicted at amino acid position 311-504 (Fig 5a). Helicase C-terminal domain profile have been predicted at aa positions 684-853 (Fig 5b). RIG-I-like receptor (RLR) C-terminal regulatory (CTR) domain profile have been predicted at amino acid site 869-998 (Fig 5c). Zinc binding domains have been predicted at amino acid positions 883, 886, 938, 941 (Fig 5d).

**Fig 5a:**
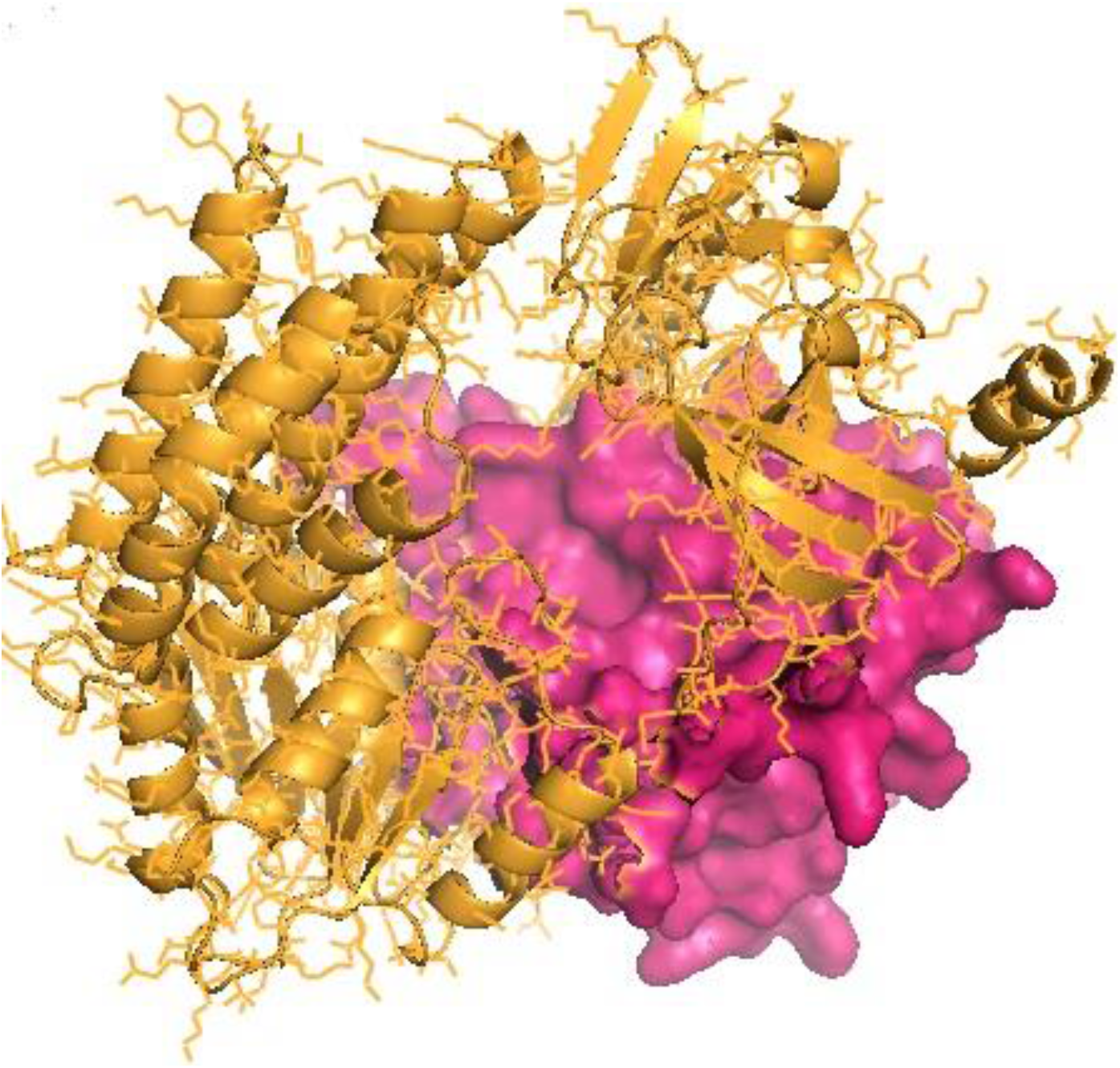
Helicase ATP binding site was predicted at amino acid position 311-504.

**Fig 5b:**
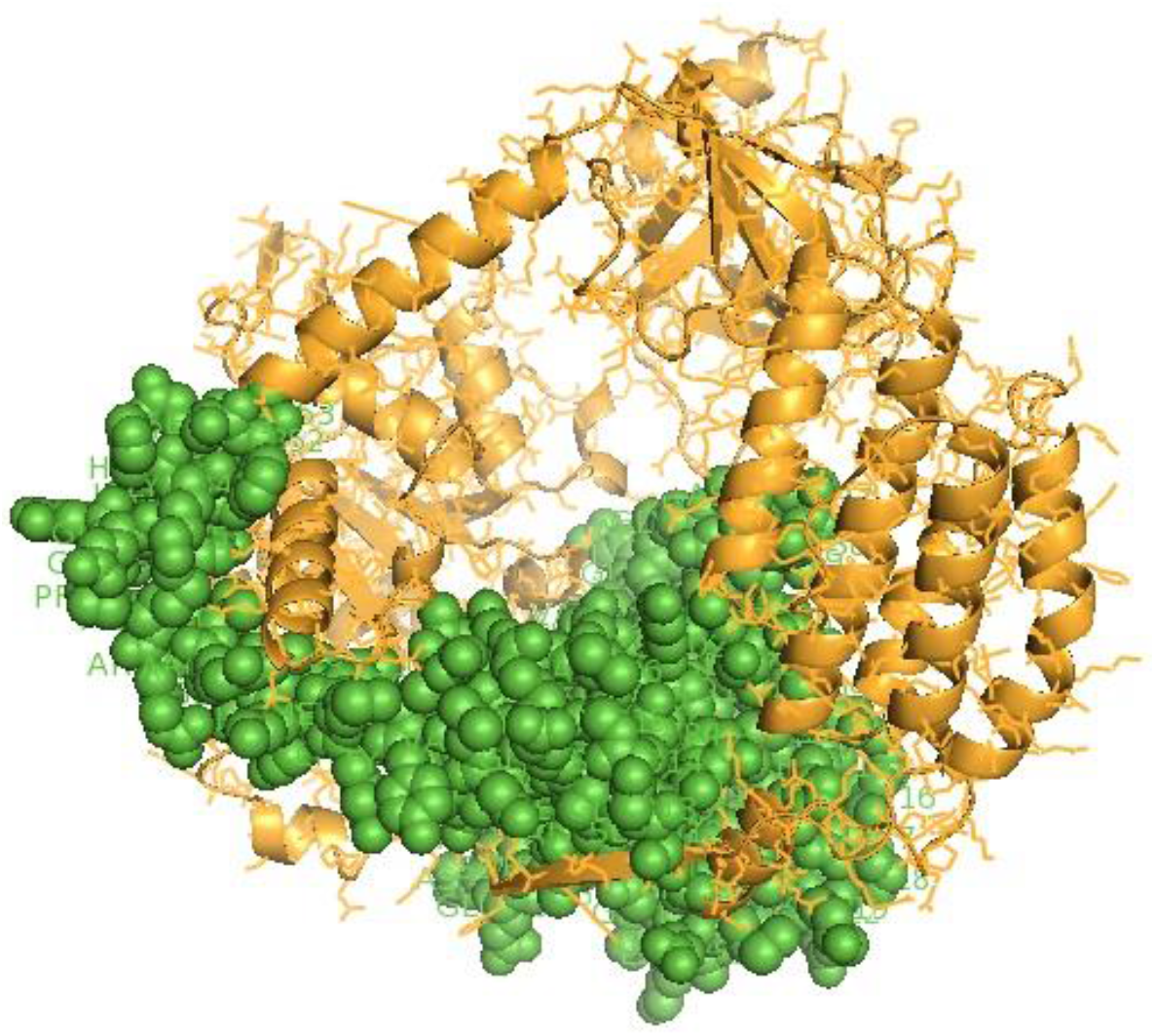
Helicase C-terminal domain profile have been predicted at aa positions 684-853.

**Fig 5c.**
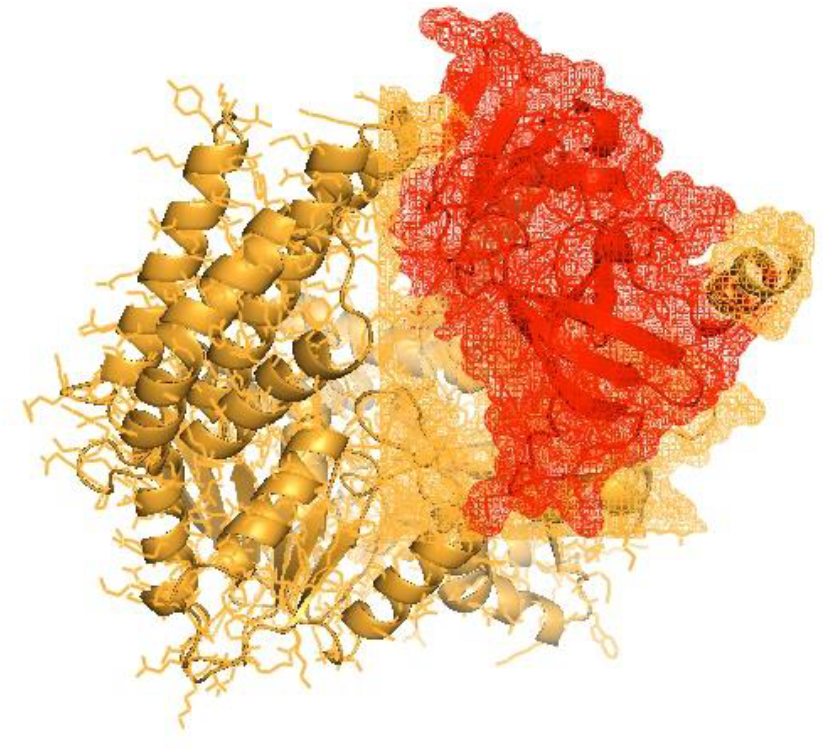
RIG-I-like receptor (RLR) C-terminal regulatory (CTR) domain profile have been predicted at amino acid site 869-998.

**Fig 5d:**
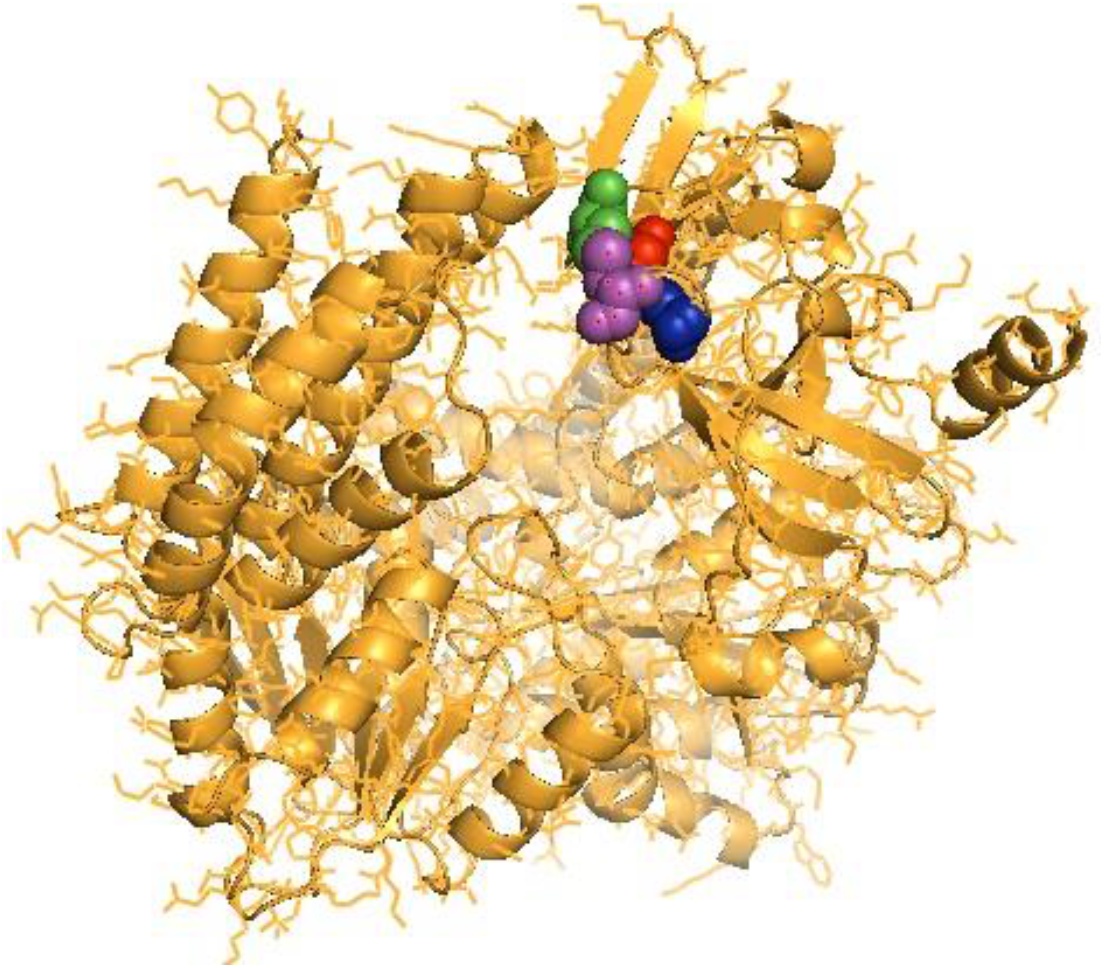
Zinc binding domains have been predicted at amino acid positions 883, 886, 938, 941.

The domain identified for duck interferon alpha reflects *Interferon alpha, beta and delta family signature* amino acid position 151-169 Fig 6.

**Fig 6:**
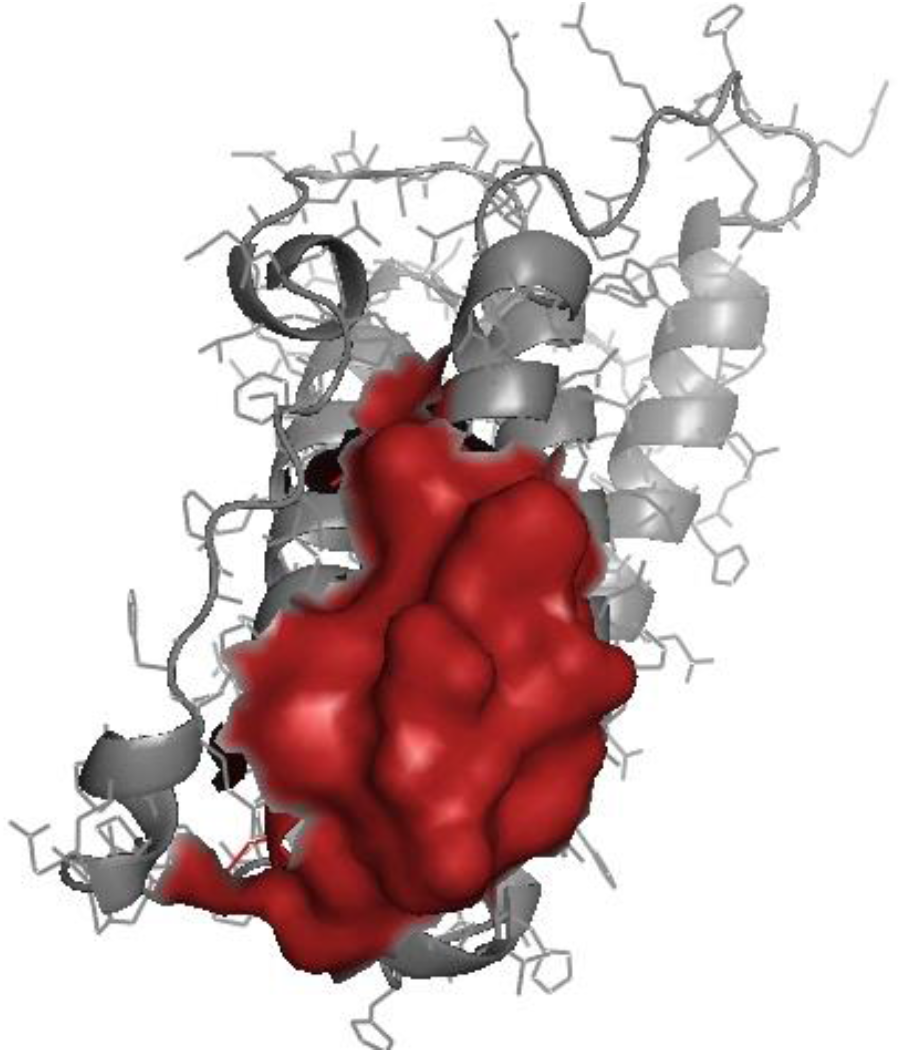
The domain identified for duck interferon alpha reflects *Interferon alpha, beta and delta family signature* amino acid position 151-169.

### Protein-protein interaction network depiction

It is important to assess the interaction between these genes to understand the immune response against Duck Plague virus interaction. RIGI (DDX58) is being represented with other genes (Fig 7a). The important genes with close interaction with RIGI are MAVS and LGP2 (DHX58). Network analysis of MDA5 and interferon alpha with other important genes is being represented in Fig 7b and Fig 7c repectively.

**Fig 7a:**
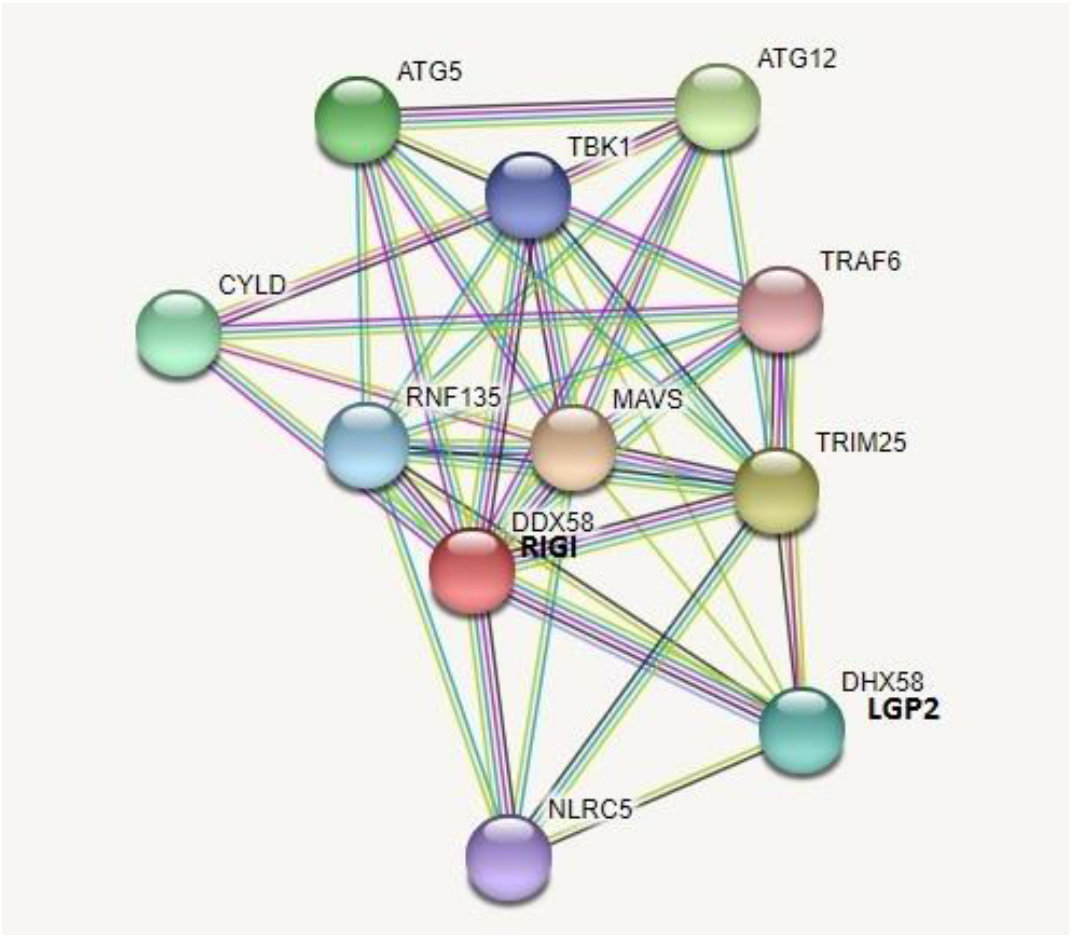
String analysis with respect to duck RIGI molecule.

**Fig 7b:**
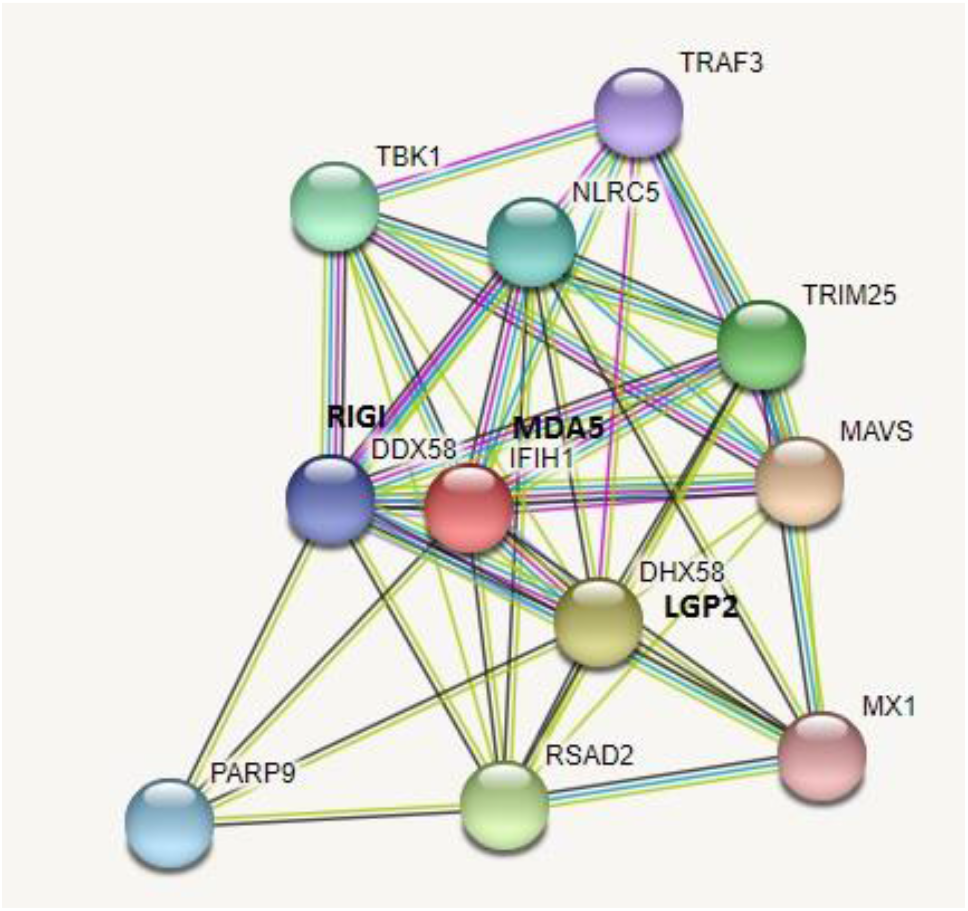
String analysis with respect to duck MDA5 molecule.

**Fig 7c:**
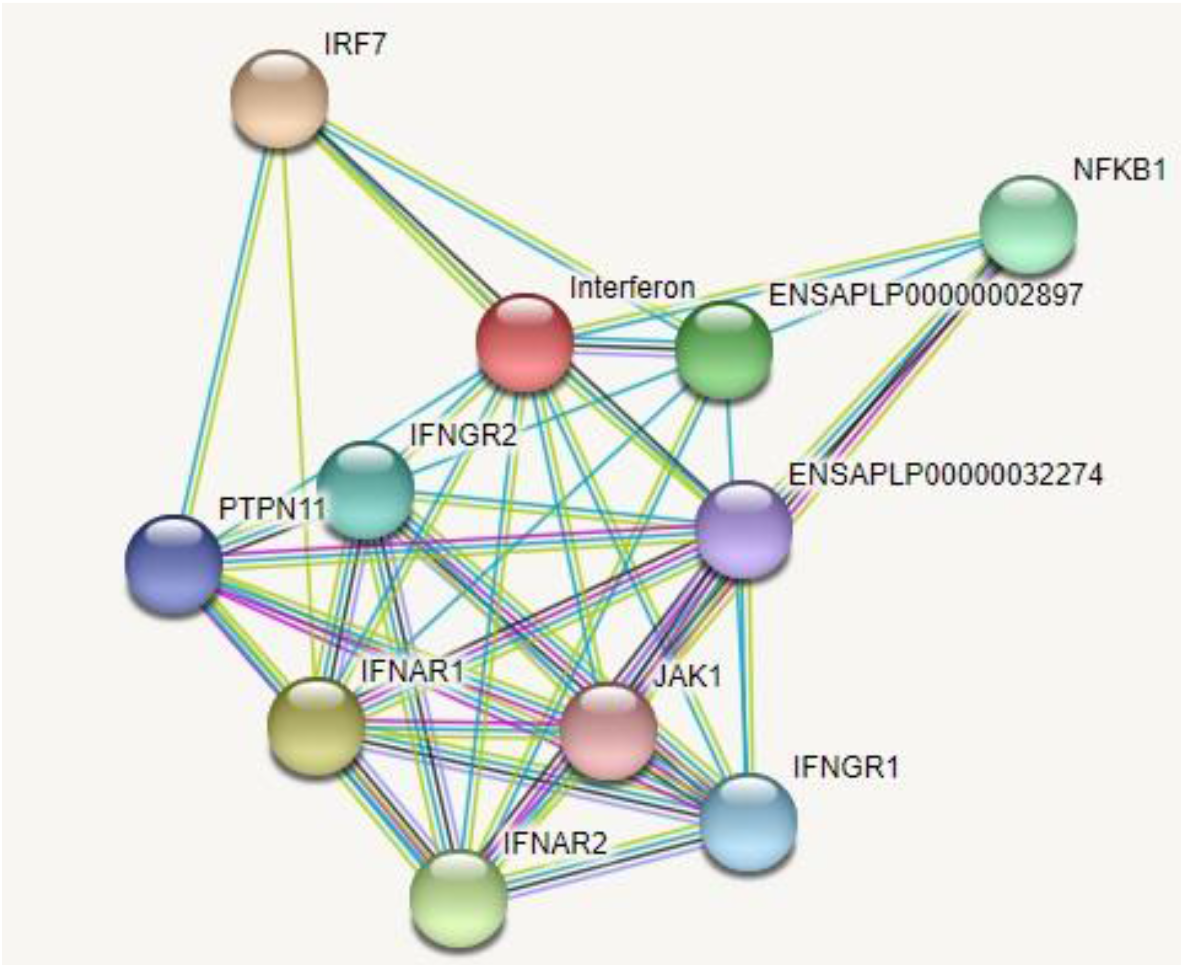
String analysis with respect to duck Interferon alpha molecule.

### *In silico* study for the detection of binding site of RIGI, MDA5, Interferon alpha, Mx, TLR7 with *Duck plague virus -Molecular docking*

We had conducted both patchdock and firedock analysis. We studied four different genes for duck plague virus (glycoprotein **MH745151**, UL30 **AFJ44192**, UL31: **HG425076**, thymidine kinase **KF214788**). Highest docking score was observed for glycoprotein, which is a structural glycoprotein for Duck plague virus (Anatid herpes virus). These studies prove the role of these genes against Duck Plague virus infection. Patchdock score has been observed to be 26066, 23136 and 19838 for RIGI, MDA5 and Interferon alpha respectively with duck plague glycoprotein.

Molecular docking was conducted for glycoprotein with genes and binding sites were predicted for these genes. We observed good binding for the genes (except Interferon). It is expected to bind since they are basically receptor molecules, whereas INFalpha is a cytokine. Molecular docking for RIGI with the duck plague viral glycoprotein indicate good score and represented in Fig 8a. (RIGI: Blue for RIGI, red for glycoprotein), whereas Fig 8b indicates the binding sites for RIGI with the glycoprotein for DP virus. The site for RIGI receptor to bind with DP viral glycoprotein ranges from Ile265(green sphere) to Arg 756 (blue sphere) as in Fig 8b. The important domains of RIGI include helicase insert domain, helicase domain interface (polypeptide binding), sites for RD interface, sites for RNA binding. DP viral glycoprotein binding site includes Ala 3 (red sphere) to Gly 269 (magenta sphere).

**Fig 8a:**
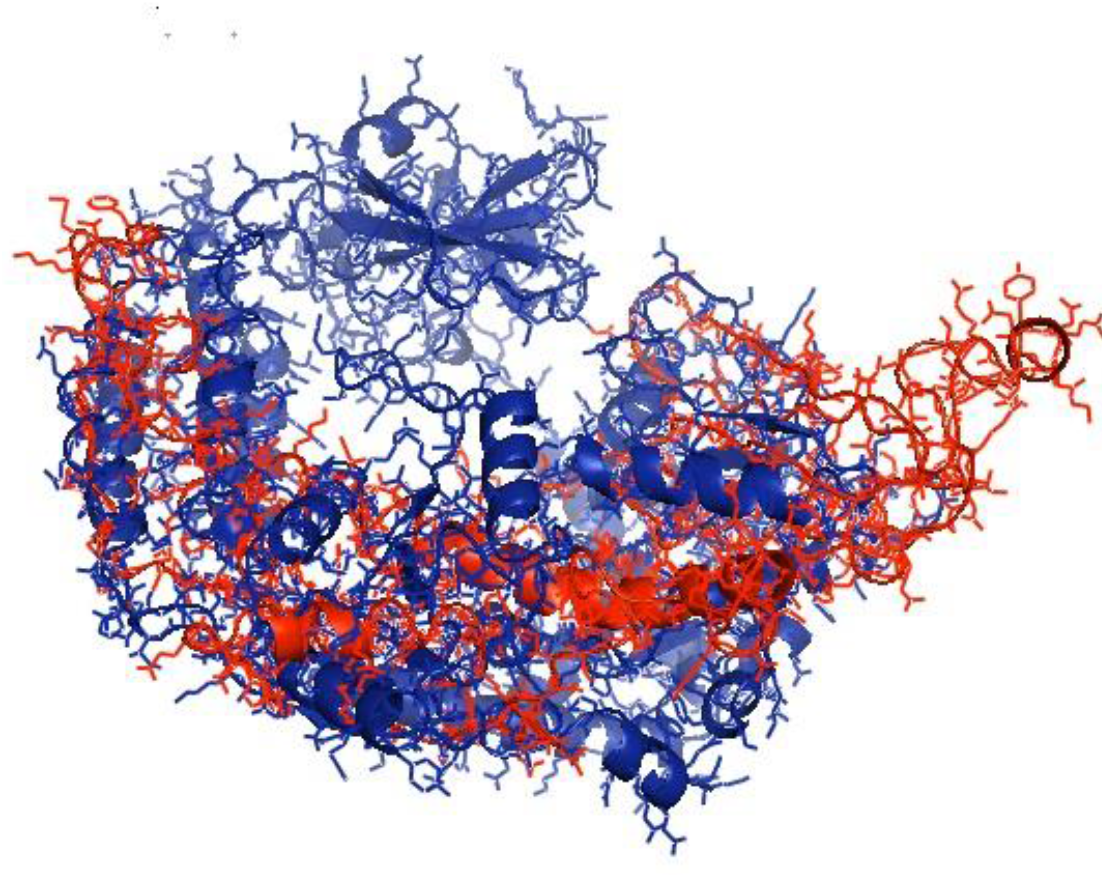
Molecular docking for RIGI with surface glycoprotein of duck plague virus.

**Fig 8b:**
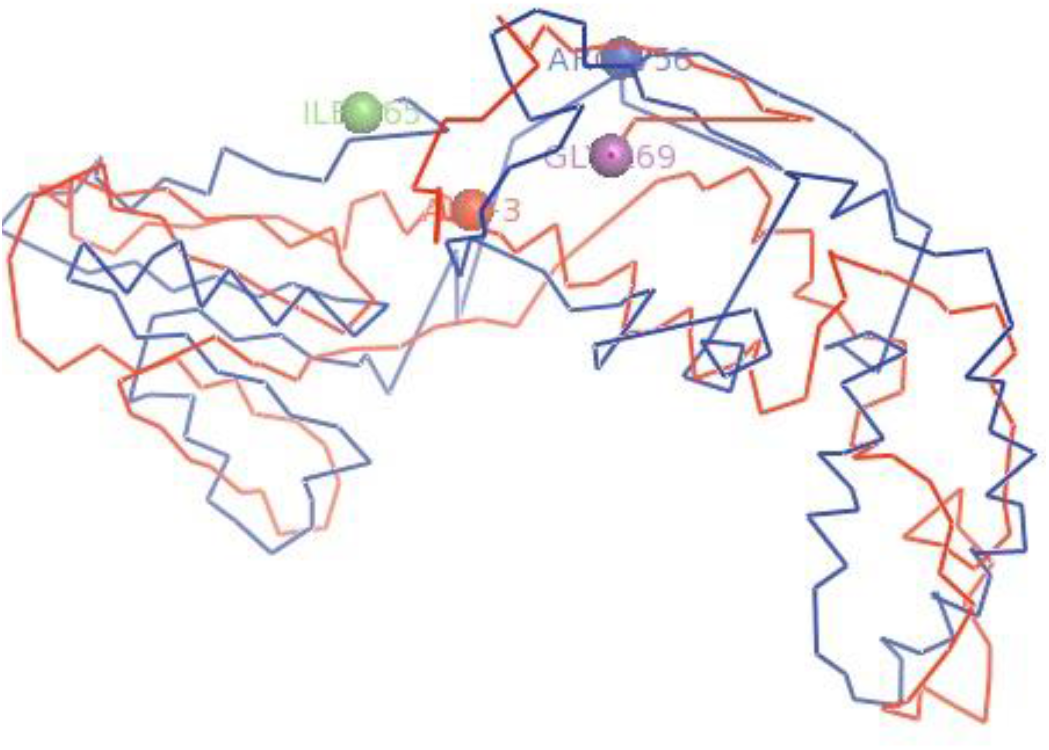
Prediction of the binding site for RIGI molecule with surface glycoprotein of duck plague virus.

Molecular docking for IFN alpha (Redfor DP viral glycoprotein) have been depicted in Fig 9a and Fig 9b represents the binding site for interferon alpha and DP virus glycoprotein. The binding site for interferon alpha with DP viral glycoprotein includes Phe29 to Thr 187.

**Fig 9a:**
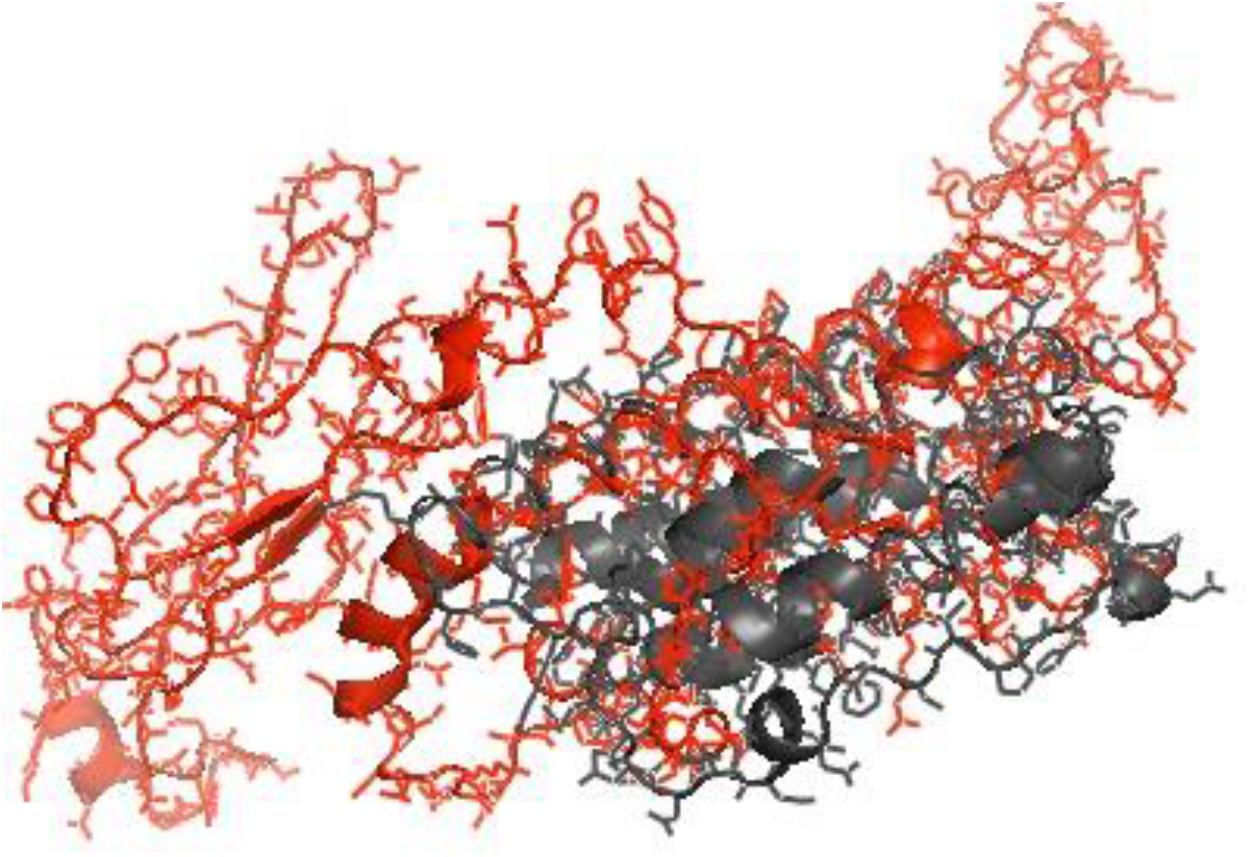
Molecular docking for MDA5 with surface glycoprotein of duck plague virus.

**Fig 9b:**
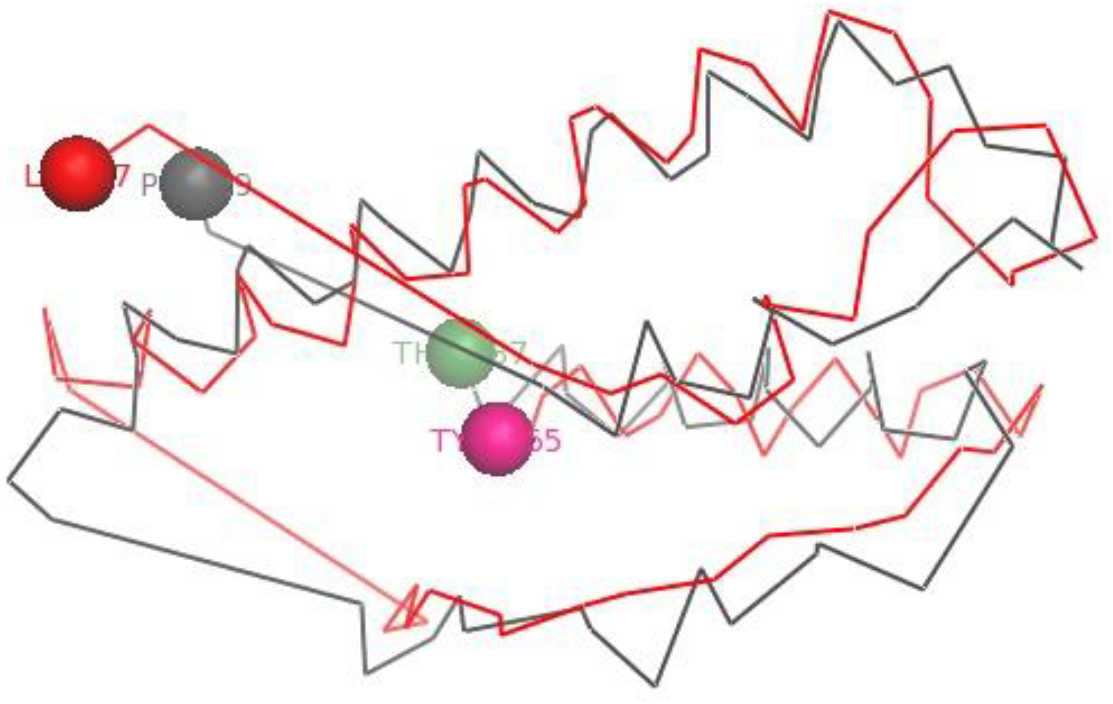
Prediction of the binding site for MDA5 molecule with surface glycoprotein of duck plague virus.

### Differential mRNA expression profiling for GALT tissues w.r.t. RIGI, MDA5, TLR7, Mx and Interferon alpha gene

Since duck plague virus mostly affects the digestive systems, we studied the expression profile for these immune response genes w.r.t. various tissues associated with gut (Fig 10). We considered liver to be one expressing most of the genes. Hence forth, we considered liver tissue for subsequent studies, since higher viral load was also detected in liver.

**Fig 10:**
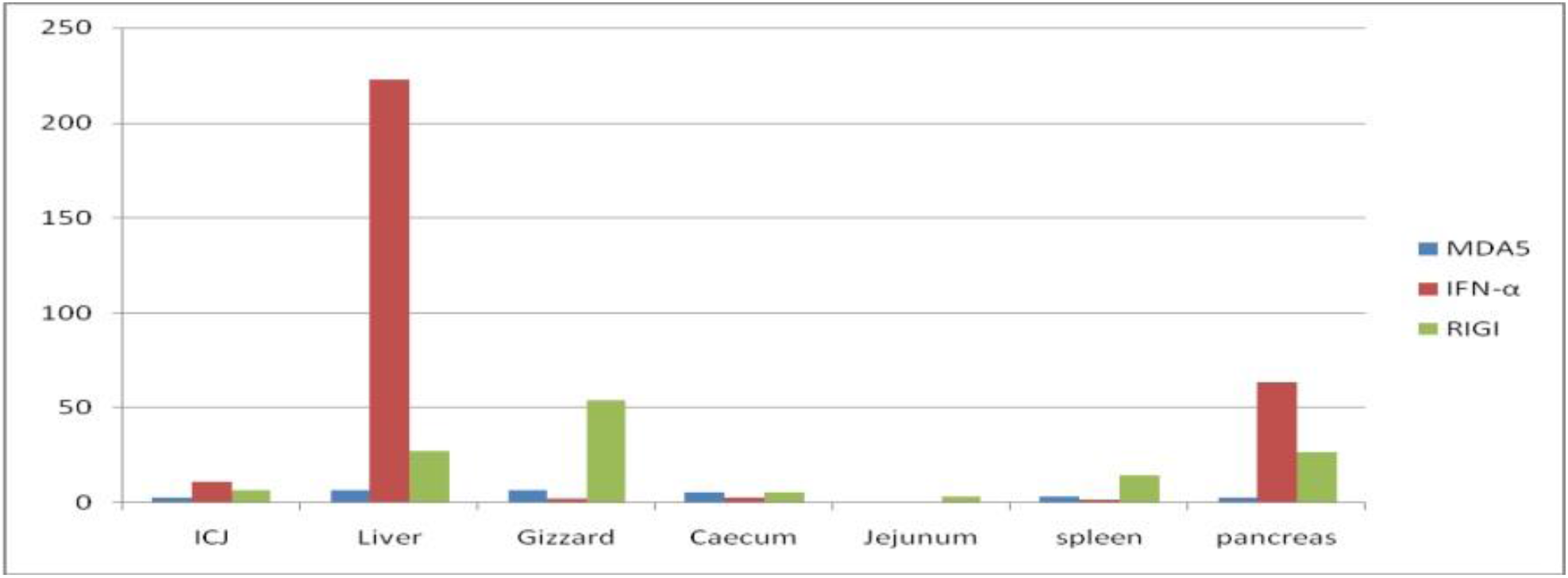
Differential mRNA expression profiling for RIGI, MDA5 and Interferon alpha with respect to GALT tissues.

### Challenge study with DP virus, symptomatic diagnosis, estimation of infective viral load at different stages of infection, PCR based detection

#### Diagnosis and confirmation for duck plague

The ducks were initially diagnosed as duck plague clinically with the help of initial symptoms. Later on confirmatory diagnosis was conducted through molecular detection of Duck plague virus in the infected samples.

#### Gross Anatomical view of the organs in duck plague infected duck

We studied different body organs from infected duck for clinical diagnosis of duck plague. The clinical symptoms have been represented in fig 11.

**Fig 11:**
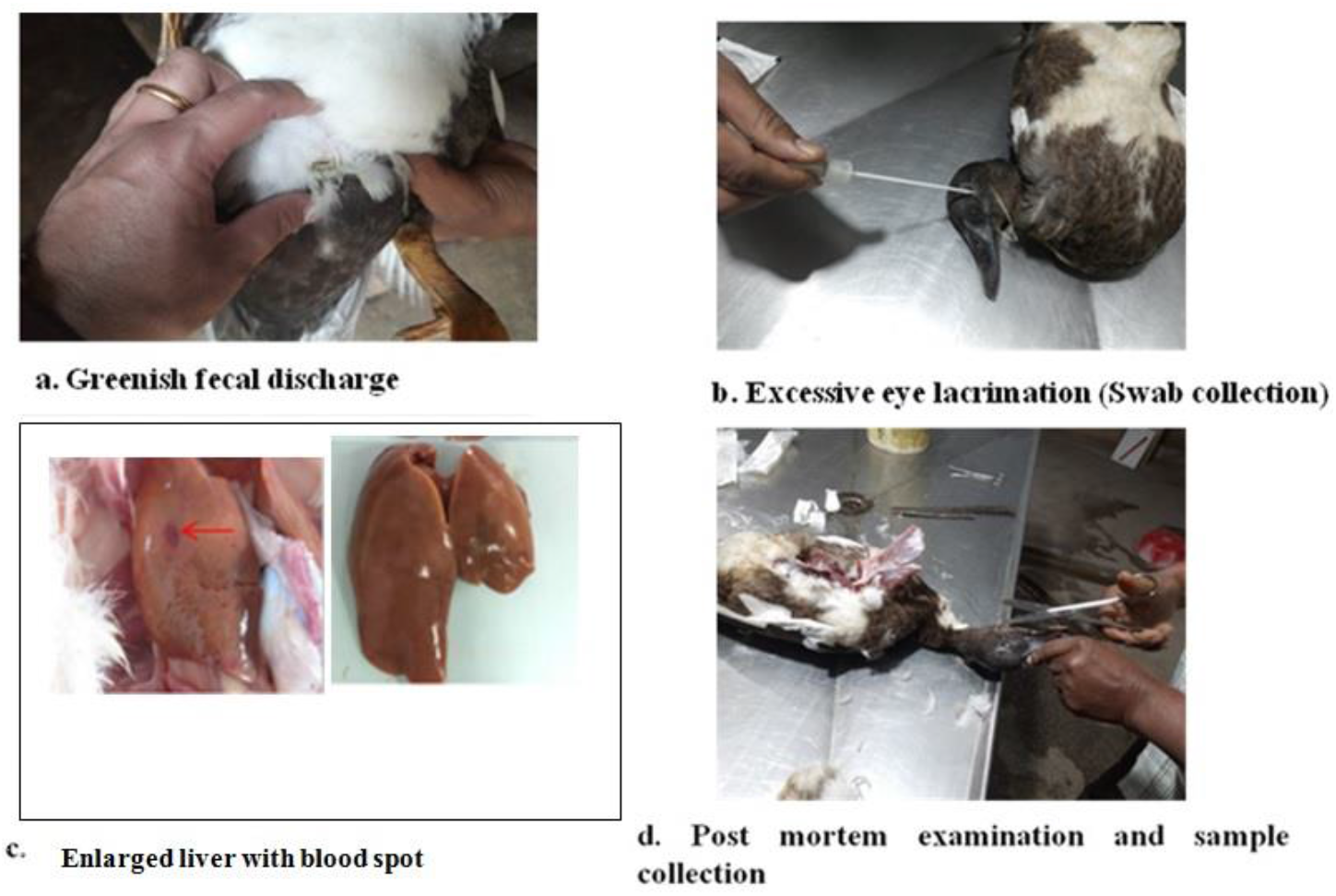
Symptomatic diagnosis of Duck plague infected duck with postmortem.

#### Confirmation of infected duck through molecular detection

In the next step, we follow confirmation of the samples with molecular PCR based technique using the following primers as recommended by OIE, Paris. It includes viral DNA isolation through isolation kit in case of DP virus and subjected to PCR. We isolate DNA since Herpes virus is a DNA virus. We detect the virus through amplification of DNA polymerase and UL44 gene (Fig 12).

**Fig 12:**
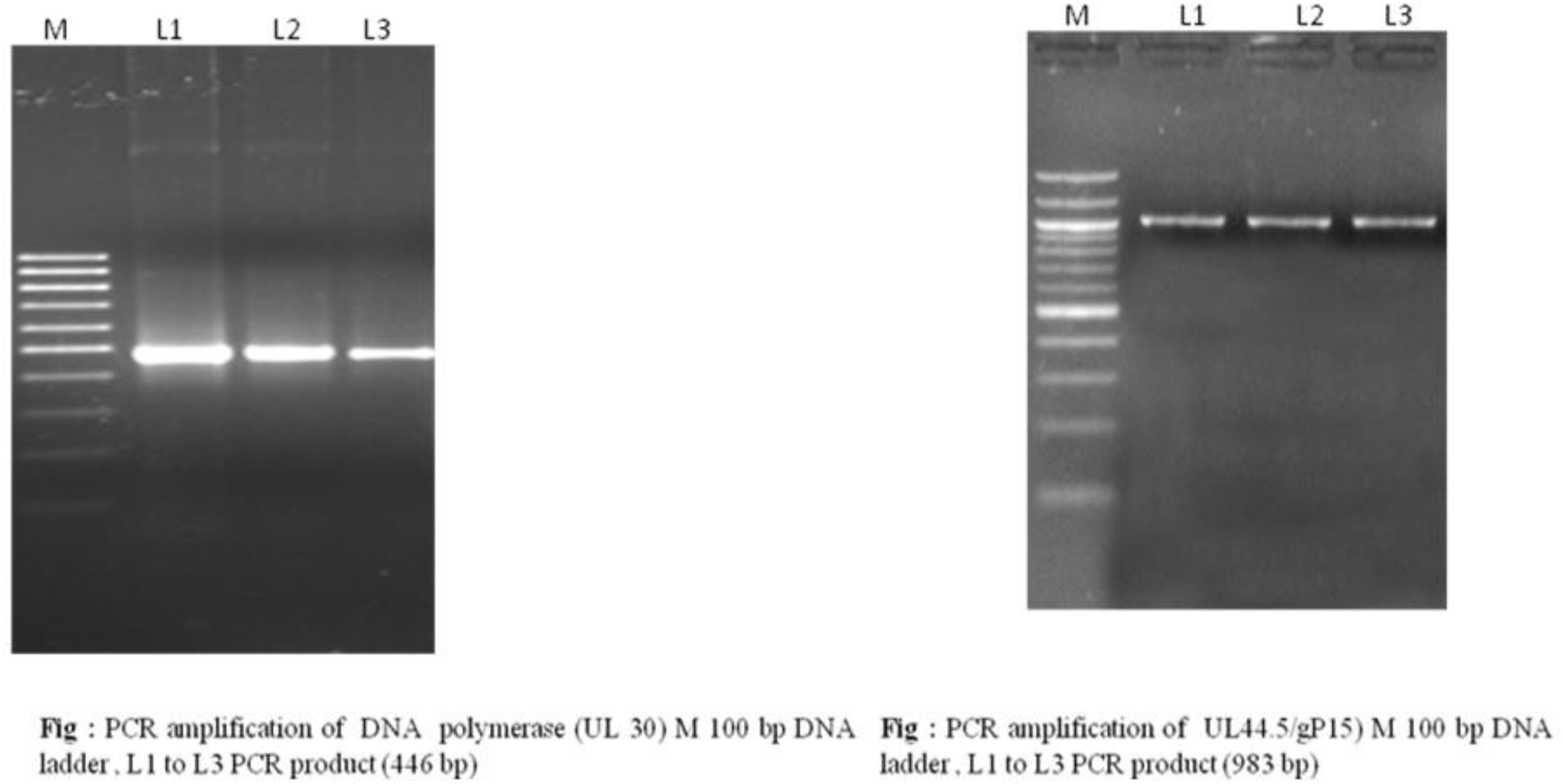
Molecular detection of duck plague infection in duck through PCR based detection of DNA polymerase and UL44 gene.

#### Assessment of dose of viral particles

We estimated the dose of viral particles through Quantitative PCR. We take the standard duck plague vaccine as a control for its amplification and quantification.

#### Estimation of biochemical parameters

The birds were examined for haematological and biochemical/ serological esimation for non-infected versus infected. The haematological and biochemical parameters are being listed in Table 4 and Table 5 respectively.

**Table 2:**
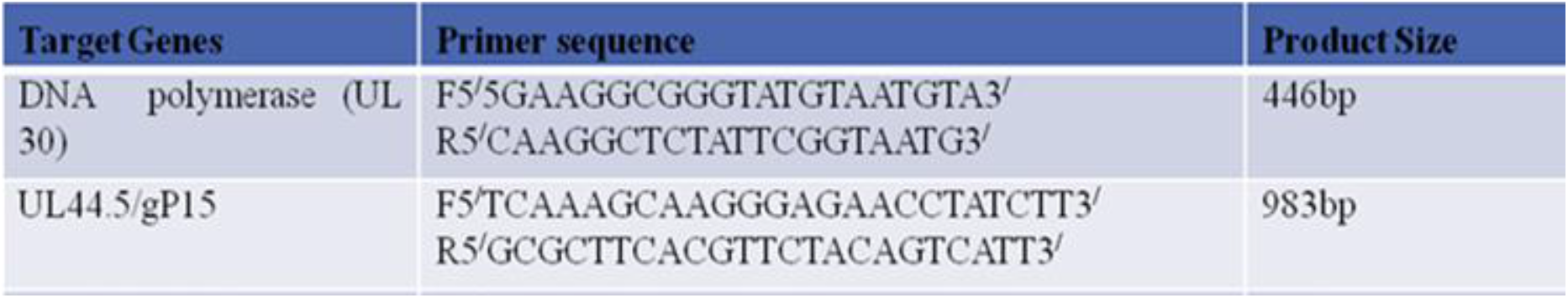
Primers used for the molecular detection of duck plague virus.

**Table 3:**
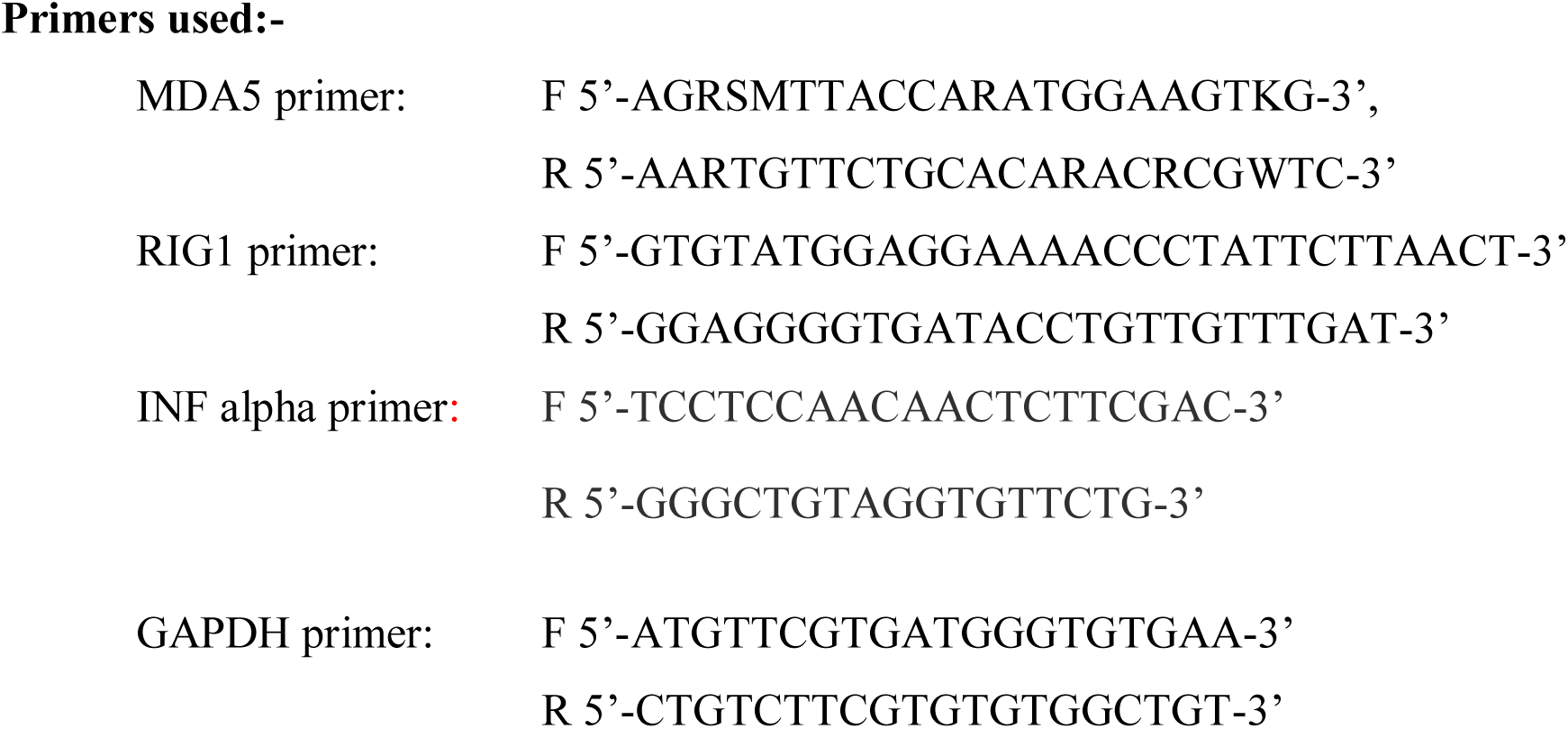
Primers used for the differential mRNA expression profiling of immune response genes of duck.

**Table 4:** Haematological profiling for healthy control ducks.

| *Parameters* | *Healthy*<br>*(N=100)* |
| --- | --- |
| **Hb (g/dl)** | 10.41 ± 0.13 |
| **PCV (per cent)** | 52.00± 0.53 |
| **TEC (× 10<sup>6</sup>/mm<sup>3</sup>)** | 2.63 ± 0.04 |
| **TLC (× 10<sup>3</sup>/mm<sup>3</sup>)** | 10.94 ± 0.22 |

**Table 5:** Biochemical parameters for infected and non-infected control ducks w.r.t. duck plague infection.

| *Parameters* | *Healthy*<br>(N=100) | *Infected*<br>(N=50) | *Level of*<br>*significance* |
| --- | --- | --- | --- |
| *Albumin (g/dl)* | *4.07<sup>a</sup> ±0.10* | *3.65<sup>b</sup> ±0.12* | *0.05* |
| *Glucose (mg/dl)* | *145.67<sup>a</sup> ±10.13* | *162.34<sup>b</sup> ±9.55* | *0.05* |
| *BUN (mg/100ml)* | *1.28<sup>a</sup> ±0.07* | *1.96<sup>b</sup> ±0.17* | *0.05* |
| *Uric Acid (mg/100 ml)* | *4.52<sup>a</sup> ±0.12* | *4.62<sup>a</sup> ±0.18* | *NS* |
| *Cholesterol (mg/100 ml)* | *120.46<sup>a</sup> ±4.73* | *119.28<sup>a</sup> ±4.37* | *NS* |
| *AST (U/L)* | *71.53<sup>a</sup> ±5.68* | *75.19<sup>b</sup> ±4.28* | *0.05* |
| *ALT (U/L)* | *21.55<sup>a</sup> ±0.17* | *22.03<sup>a</sup> ±0.28* | *NS* |

#### Differential mRNA expression profiling for Immune rsponse genes in infected and healthy ducks

Upregulation was observed for genes RIGI and Interferon alpha in infected duck w.r.t. healthy control (Fig 13). We observed downregulation for MDA5 gene expression in infected birds in comparison to non-infected control.

**Fig 13:**
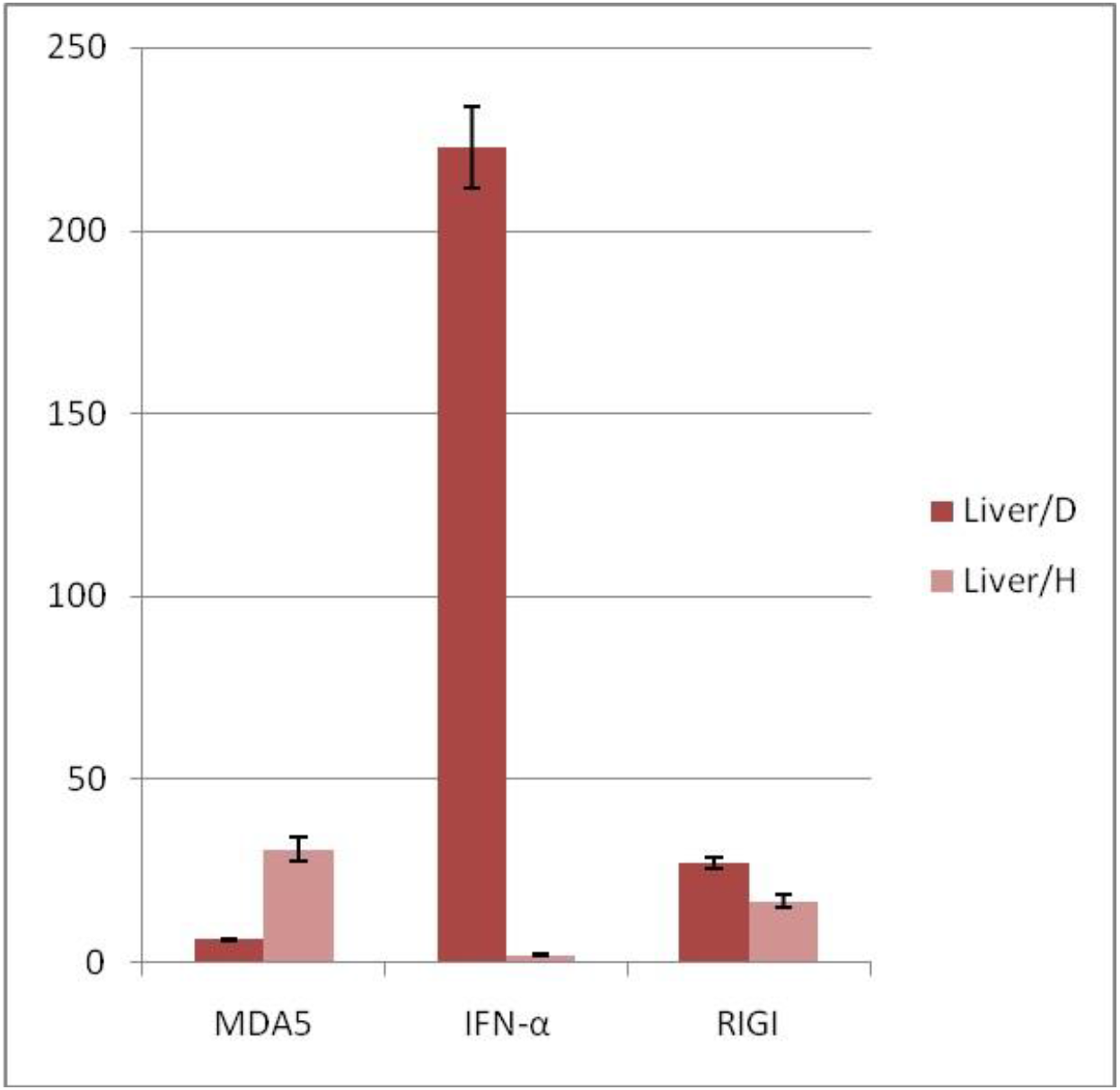
Differential mRNA expression profiling for RIGI, MDA5 and interferon alpha genes in liver tissue in infected and healthy control w.r.t.duck plague infection.

#### Histological section and immunohistochemistry

Light-microscopic images of Mayer’s Hematoxylin-stained optical sections of liver from the control healthy subjects A have been depicted in Fig 14a, and that of infected samples in Fig 14b. Microscopic visualization of Kupfer cells by the Cavalieri-physical disector combination method is being represented in Fig 15.

**Fig 14a:**
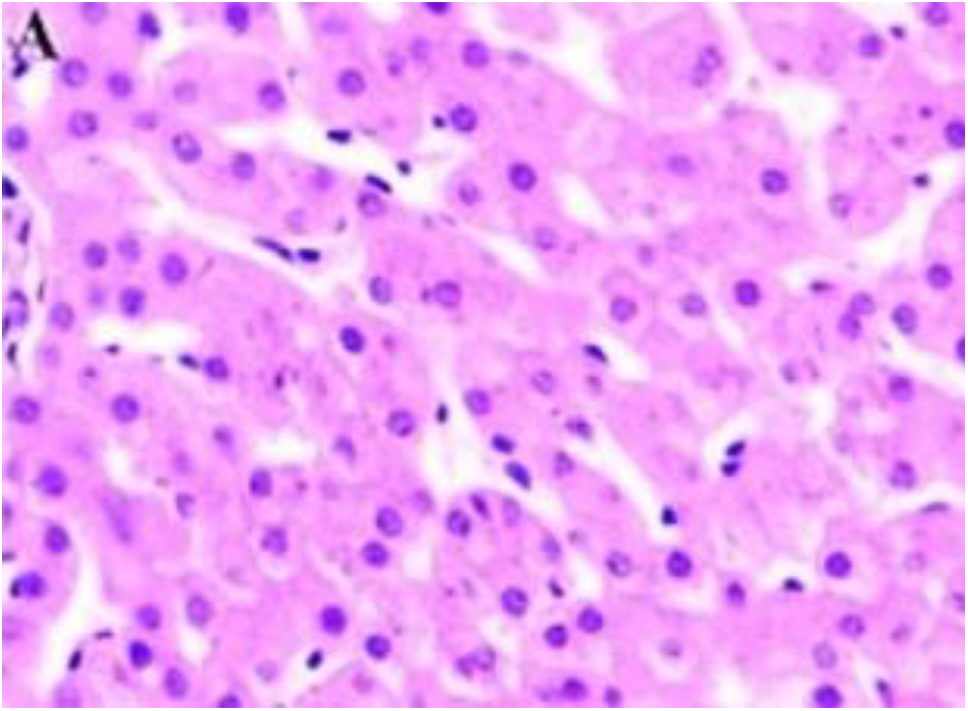
Histological section for liver tissue of healthy control duck.

**Fig 14b:**
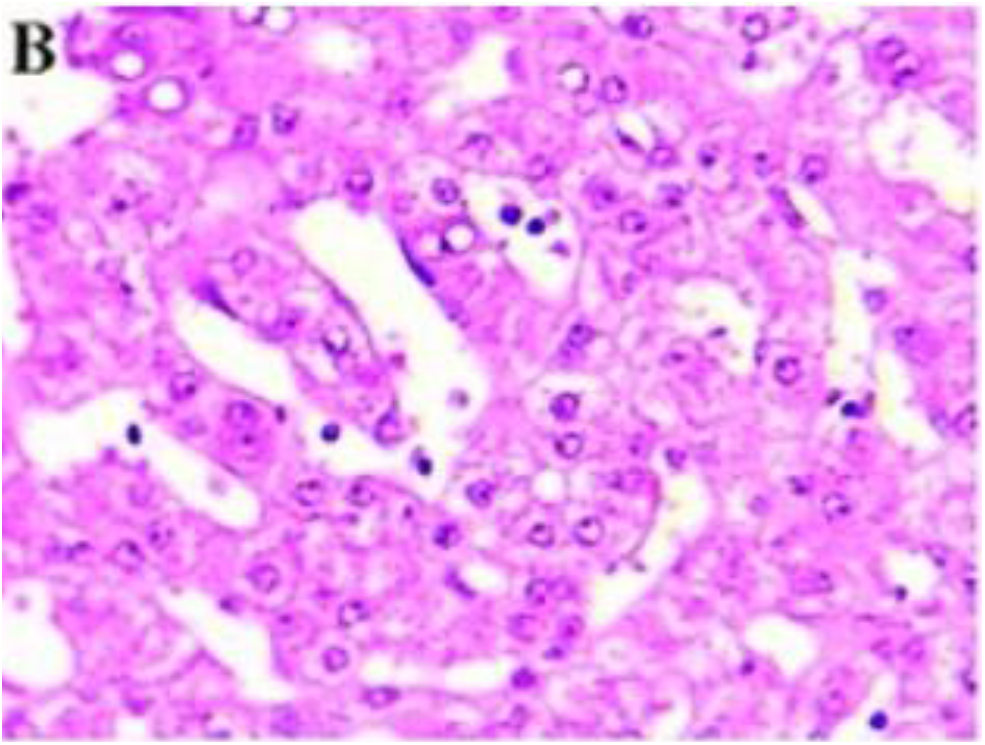
Histological section for liver tissue of duck plague infected duck.

**Fig 15:**
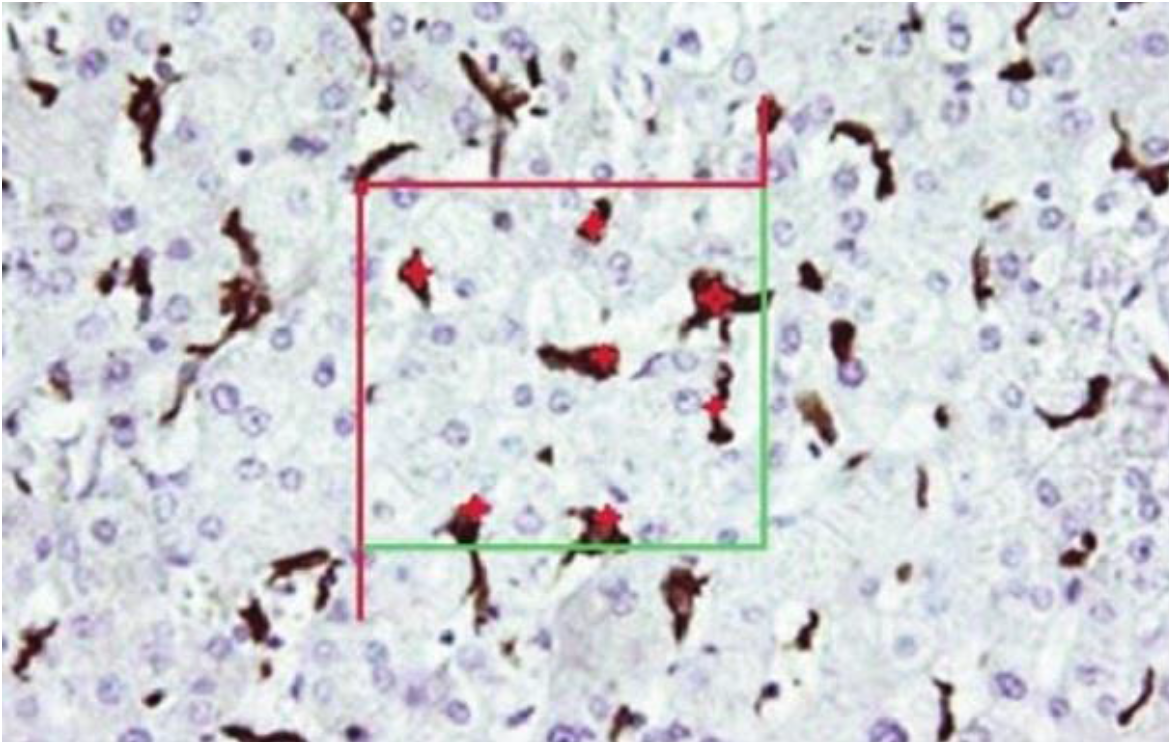
Microscopic visualization of Kupfer cells by the Cavalieri-physical disector combination method.

Immunohistochemistry for the expression of RIGI protein and MDA5 in the healthy have been represented in Fig 16 and Fig 17 respectively.

**Fig 16:**
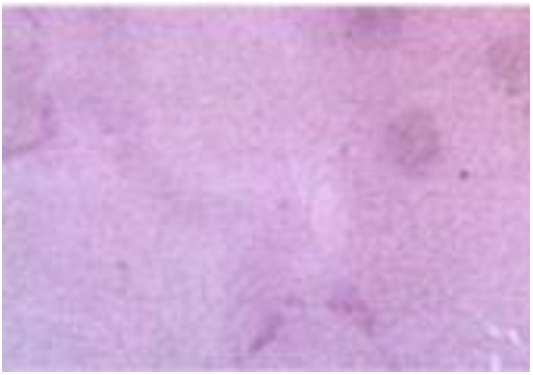
Immunohistochemistry for RIGI in healthy duck.

**Fig 17:**
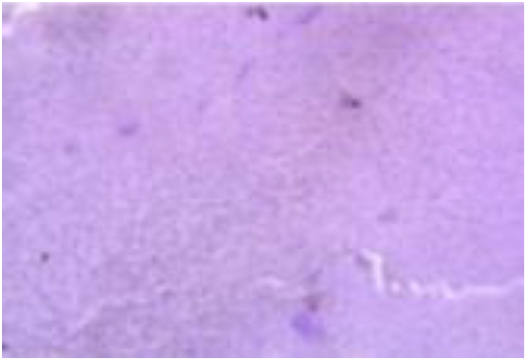
Immunohistochemistry for MDA5 in healthy duck.

## Discussion

Duck plague is caused by anatid herpes virus 1 which is a linear double stranded DNA virus. We studied the gene sequences for the four important genes of duck plague virus (UL30, UL31, thymidine kinase and glycoprotein) for *in silico* molecular docking analysis. These genes were sequenced earlier by researchers from our university. UL31-It helps in late phase replication (https://www.ncbi.nlm.nih.gov/pmc/articles/PMC2661054/). UL30 is involved in DNA replication and nucleotide metabolism in DPV (https://www.ncbi.nlm.nih.gov/pmc/articles/PMC8057227).

Duck plague virus glycoprotein (DPV gI) plays major roles in viral cell-cell spread and viral final envelopment process. DPV gI protein played vital roles in viral cell-to-cell spread and virion final envelopment. It also provided a communication that DPV gI may affect the gE TGN location to influence virion final envelopment (https://www.sciencedirect.com/science/article/pii/S003257912030612X). The thymidine kinase (TK), as the main enzyme in replication of DPV, is probably expressed at early times after infection, which caused its quick immune recognition after the infection. TK is a prior target gene for constructing the gene-deleted vaccine (https://virologyj.biomedcentral.com/articles/10.1186/1743-422X-7-77). We observed highest docking score for surface glycoprotein. This is quite obvious since for binding with any receptor, the surface glycoprotein is of utmost importance. Glycoproteins E (gE) is not essential for the extracellular entry of HSV and PRV but plays a pivotal role in directing virion spreading from cell-to-cell in polarized cells, such as epithelial cells and neuronal cells. The gE cytoplasmic (CT) domain aid the virus to carry out cell-to-cell spreading by trafficking virions through the trans-Golgi network (TGN) to the lateral surfaces of polarized cells. The extracellular (ET) domains of glycoprotein could bind ligands at cell junctions, thereby promoting the virus entry into adjacent cells. Furthermore, gE plays essential roles in the replication of specific alphaherpesviruses, such as MDV-1 and VZV. Duck plague viral surafce glycoprotein is also responsible for providing essential but redundant functions during the acquisition of the final virion envelope. In HSV-1, gE together with gD affects the production of enveloped virus particles. DPV gE played a significant role in the final envelopment process and provided new information for the study of the molecular mechanism of DPV (Liu et al., 2020).Since this is the first report of molecular docking with DP virus, comparison was not possible. It is interesting to note that DPV have CPG motif in their genome.

In this study, we could observe better expression of these immune response genes (RIGI, MDA5 and Interferon alpha) in liver in comparison to the other organs of gut under study, namely ileocaecal junction (caecal tonsil), gizzard, caecum, small intestine, spleen and pancreas. Highest expression of interferon alpha was observed in liver followed by pancreas, then ileocaecal junction or caecal tonsil. A trend was observed for lower level of expression of MDA5 gene in GALT. Since liver expresses these immune response genes sinificantly, we had considered liver as GALT for futher subsequent studies. Liver possess kupfer cells, which are the major macrophages found in the sinusoids of liver. They constitute for 80-90 percent of all the macrophages in the entire body. We could visualize the kupfer cells in healthy subjects. GALT present in pancreas is islet of langerhans. We observe higher expression of RIGI in proventriculus and ventriculus (gizzard), which may be due to the presence of diffuse lymphoid tissue in the epithelium or pyloric tonsil (Casteleyn et al., 2010) tissues (gut associated lymphoid tissues) differ in different organs. We observed the highest expression of RIGI in mocosal layer of gizzard, followed by liver and pancreas.

The current study reveals better expression profile of RIG1 in infected duck compared to non-infected control, which implies a role of RIG1 in binding with duck plague virus and neutralizing it with help of different cytokines. Higher binding score of RIGI with DP virus surface glycoprotein have reported through *in silico* molecular docking studies. Role of RIG1 with double stranded DNA has already been documented (UniProtKB-095786, DDX58_HUMAN). It was also found that MDA5 mRNA epression in healthy liver tissue is more than duck plague infected samples, where as RIG1 shows a different result. In the current study we observed a very high expression potential for Interferon alpha in infected samples, however through in silico studies, the binding score was observed to be less.

Since, this is the first report of the study of expression profiling of liver with respect to duck plague infection, the comparison as such was not possible. In an earlier single study to assess host immunity in response to duck plague infection, similar findings of upregulation of RIGI and interferon alpha, whereas downregulation of MDA5 were observed in spleen with duck plague viral infection (Li et al.,2016). However they had observed a different finding with brain. It is evident that microglial cells are important in immune cells in brain, whereas lymphocytes are important in spleen. Hence it is evident that tissue tropism is an important criteria.Since the major lymphoid tissue in liver is also macrophages, we also observed a similar finding as in spleen.

Innate immune system constitute of some genes which are commonly known as immune response gene are classified as the type of receptors such as RLR, TLR etc. Both RIG1 and MDA5 belong from the same family and found in cytoplasm (Wei et al.). It was observed by Zou et al. that vertebrate species having conserved sequences of MDA5 and sometime expressed in a low level but it may be induced by viral challenge.

In our earlier study, we observed that RIGI was better expressed in avian influenza (of H5N1 strain) virus infected ducks in comparison to healthy control (Pal et al., 2021). It is to be noted that avian influenza virus is a RNA virus, whereas duck plague virus is a DNA virus. Through molecular docking analysis, it was observed in both the cases that RIGI effectively binds with both H and N antigen of avian influenza virus as well as surface glycoprotein of duck plague virus with good score. After initial binding, it is important to analyze the mode of pathogenesis in either cases. Duck plague virus is expected to be similar to alpha herpes virus of human, since both belong to alpha herpes virus subfamily (Liu et al., 2020, Zhao et al., 2021).However, till date we could not explore any report for zoonosis, which need to be explored in future.

Herpes virus affecting human being is a similar DNA virus within the Herpesviridae family as in duck plague virus. Similarly hepatitis B virus is also a DNA virus. It is essential to study the molecular pathogenesis of duck plague virus and how these RIGI and MDA5 acts.There may be two basic mode of action after binding, either they directly destroy the DNA (genetic material) of the virus. Otherwise they may act on the transcribed RNA coded by the DNA virus as DP virus. From the available literature, it is evident that they basically act on ssRNA, dsRNA and dsDNA viruses (Dias Junior et al., 2019).

There exists some basic differences in the mechanism of RIGI and MDA5. The signaling pathway for RIGI/MDA5 depicts that the initial step involves recognition of PAMP dsRNA, leading to the interaction with MAVS. After the activation of MAVS by RIG-I/MDA5, a molecular cascade initiates the interaction of IKKε and TBK1. The next step is the phosphorylation of the transcription factors IRF3 and IRF7, ensure to translocate the phosphorylated p-IRF3 and p-IRF7 into the nucleus. These dimerize and bind to transcription factor binding sites of the IFNα and IFNβ genes for their activation of their transcriptions. Expression and exportation of these genes into the cellular milieu activate the IFN1 signaling cascade in an autocrine or paracrine fashion for the expression of hundreds of interferon stimulated genes (ISGs) and inflammatory genes for confering antiviral resistance. RIG-I and MDA5 also activate the NF-κB pathway. RIG-I appears to act upstream of the canonical pathway, resulting in the translocation of the two functional NF-κB units (p50 and p65) into the nucleus. On the other hand, MDA5 is reported to affect NF-κB expression independently from this pathway (Dias Junior et al., 2018).

It has been observed that everal HSV-1 proteins could down-regulate the RLR signaling pathway via targeting multiple proteins involved in this pathway with distinct mechanisms ( Su et al.,2016). US11, an RNA binding tegument protein of HSV-1 [Johnsn et al., 1986], interacts with endogenous RIG-I and MDA5 via its carboxyl-terminal amino acids in an RNA independent manner, and subsequently impedes the formation of RIG-I/MAVS and MDA5/MAVS complexes, resulting in reduced production of IFN-β [Zing et al., 2012].

Hence, it is evident from our study that MDA5 could bind with glycoprotein of the DP virus, but could not destroy it. Expression analysis reveals that MDA5 is not trigerred in infective cases in liver.

## Conclusion

We could characterize the gene for RIGI, MDA5 and INF alpha protein from liver tissue of duck origin. We observed good binding ability of RIGI and MDA5 with the surface glycoprotein of Duck plague virus, a dsDNA virus, belong alphaherpesvirus sub family and Alphaherpesviridae family.Differential mRNA expression profiling reveals better expression of RIGI and pronounced expression of interferon alpha in duck plague virus challenged samples. This study proves positive role of RIGI and interferon alpha as immune response genes against duck plague viral infection. This information may be helful for production of duck with the inherent resistance against duck plague virus infection through suitable biotechnological approaches as gene editing. It may also open up the scope to study host immunity against herpes virus in animal model.

## Acknowledgement

The authors are thankful to Department of Biotechnology, Ministry of Science and Technology, Govt. of India (Grant number BT/PR24310/NER/95/649/2017) and Department of Science and Technology, Govt. of India (Grant no. EMR/2016/003554) for providing the financial support. The technical and financial support by Vice-Chancellor, West Bengal University of Animal and Fishery Sciences is duly acknowledged. Thanks to Director, AH & VS, Animal Resource Development Department, Govt. of West Bengal.

